# Spatially Distinct Macrophage Subsets Drive Myofibroblast Heterogeneity and Maladaptive Fibrosis in Lupus Nephritis

**DOI:** 10.64898/2026.04.27.719870

**Authors:** Chirag Raparia, Paul Hoover, Junting Ai, Marcus Clark, Sujal Shah, Accelerating Medicines Partnership (AMP) RA/SLE Network, Betty Diamond, Nir Hacohen, Arnon Arazi, Anne Davidson

## Abstract

**Objectives:** Lupus nephritis (LN) is a severe complication of systemic lupus erythematosus (SLE), leading to progressive renal fibrosis and functional decline. Understanding the interplay between immune cells and stromal cells is needed to develop effective therapeutic strategies. Here, we investigated the landscape of macrophage-fibroblast interactions in human LN and validated these findings in mouse models.

**Methods:** We characterized distinct fibroblast subsets and their interactions with renal macrophages using single-cell RNA sequencing (scRNAseq) of 156 human LN biopsies and 30 healthy controls from the AMP-SLE cohort, and spatial transcriptomics of biopsies from 6 LN patients. *In vitro* co-culture studies using mouse models were performed to further define functional consequences of these interactions.

**Results:** We identified two myofibroblast subsets: a pro-inflammatory subset (Myofib1) enriched in the tubulointerstitium, and a fibrotic/remodeling subset (Myofib2) in glomeruli, both correlating with the histologic chronicity index. Spatial transcriptomics revealed different colocalization patterns, with Myofib1 interacting with activated resident macrophage (RM) subsets and Myofib2 with glomerular infiltrating disease-associated macrophages. *In vitro* co-culture studies demonstrated that nephritic RMs promote a pro-inflammatory, remodeling fibroblast phenotype that impairs wound healing and drives a Myofib1-like gene program, whereas disease-associated macrophages generated profibrotic fibroblasts with dysregulated reparative capacity. Cell-cell communication analyses identified key ligand-receptor interactions mediating this crosstalk, including Spp1/integrins, Sema4/PlexinB, and NAMPT/INSR.

**Conclusions:** Our data reveal a spatially and functionally heterogeneous landscape of macrophage-fibroblast crosstalk in LN. These findings advance our understanding of renal fibrogenesis in LN, highlighting specific fibro-inflammatory circuits that may represent therapeutic targets to prevent chronic renal damage.

## INTRODUCTION

Lupus nephritis (LN) is a severe complication of systemic lupus erythematosus (SLE) leading to chronic renal failure in 10-15% of patients ^1^. Kidney fibrosis in LN is a maladaptive injury response characterized by glomerulosclerosis and interstitial accumulation of injured proximal tubular cells, endothelial cells, myofibroblasts, extracellular matrix, and immune cells in the fibrogenic niche ^2, 3^. The chronicity index, comprising glomerulosclerosis, fibrous crescents, tubular atrophy and interstitial fibrosis, quantifies fibrotic changes in LN biopsies and associates with poor treatment responses and renal functional decline ^4^.

In response to injury or inflammation, macrophage-fibroblast interactions promote differentiation of fibroblasts into heterogeneous states that are conserved across species, including myofibroblasts that are the major source of renal collagen and other matrix components ^5, 6, 7^. During physiological repair myofibroblasts die or are deactivated, but their persistence in chronic disease is a key driver of irreversible organ injury ^8, 9^.

Renal macrophages manifest diverse phenotypes that can promote renal injury, repair/resolution, or fibrosis ^10, 11^. In previous scRNAseq studies of four mouse LN models we identified multiple renal macrophage subsets that occupy different tissue niches. Two of these, classical 2 (C2) monocytes and resident macrophage cluster 0 (RM0) arise exclusively in kidneys of nephritic mice and appear to differentiate *in situ* from infiltrating Classical 1 monocytes and homeostatic resident macrophages (RM1) respectively ^12^. C2 and RM0 macrophages express a damage-associated gene expression profile ^12, 13^ (including Trem2, CD9, CD63, Spp1, Gpnmb, and Fabp5) that has been observed in other fibrotic organs ^11, 14^. In human LN kidneys, C2 monocytes localize to the glomeruli, and their frequency correlates with the histologic activity score, whereas RM0 occupy the interstitium and correlate with chronicity ^12^. However, their functional roles in fibrosis vs. repair remain unclear.

Here we mapped the spatial relationships of renal macrophage subsets with fibroblasts in human LN kidneys and performed co-culture studies of fibroblast differentiation and function using isolated mouse RMs and induced C2-like monocytes.

## METHODS

These are included in the Supplementary Data.

## RESULTS

To characterize the diversity of fibroblasts in LN kidneys we analyzed scRNASeq data from the Accelerating Medicines Partnership SLE (AMP-SLE) cohort comprising 156 LN patients and 30 healthy controls (**Supplementary Table 1**) ^15^. Coarse clustering of 16,187 stromal cells yielded separate clusters of mesangial cells, fibroblasts, myofibroblasts, endothelial cells and 2 subclusters of vascular smooth muscle cells based on expression of Calponin-1 (Cnn1) (**Supplementary Figure 1A, B**). High resolution subclustering identified 4 fibroblast clusters, 2 myofibroblast clusters, 10 vascular smooth muscle cell (VSMC) clusters and 5 endothelial cell clusters (**Figure 1A, Supplementary Figure 1C-E, Supplementary Table 2**). Compared with healthy controls, LN patients showed an increased frequency of fibroblasts, myofibroblasts and mesangial cells and a decrease in Cnn1^+^ pericytes (**Figure 1B**). Covarying neighborhood analysis (CNA) revealed no significant correlations between stromal subcluster frequency and the histologic activity index (**Supplementary Figures 2A-F**). By contrast, the chronicity index correlated with a decreased frequency of mesangial cells and an increased frequency of myofibroblasts and Cnn^-^ VSMC/Pericyte1 (**Figure 1D**). Correlations with myofibroblasts and pericytes were also present for the tubular atrophy and interstitial fibrosis subscores (**Figure 1D, 1E**). Mesangial cell loss correlated with the glomerulosclerosis score (**Figure 1F**) but not with cellular or fibrous crescents (**Supplementary Figure 2G, 2H**).

**Figure 1:**
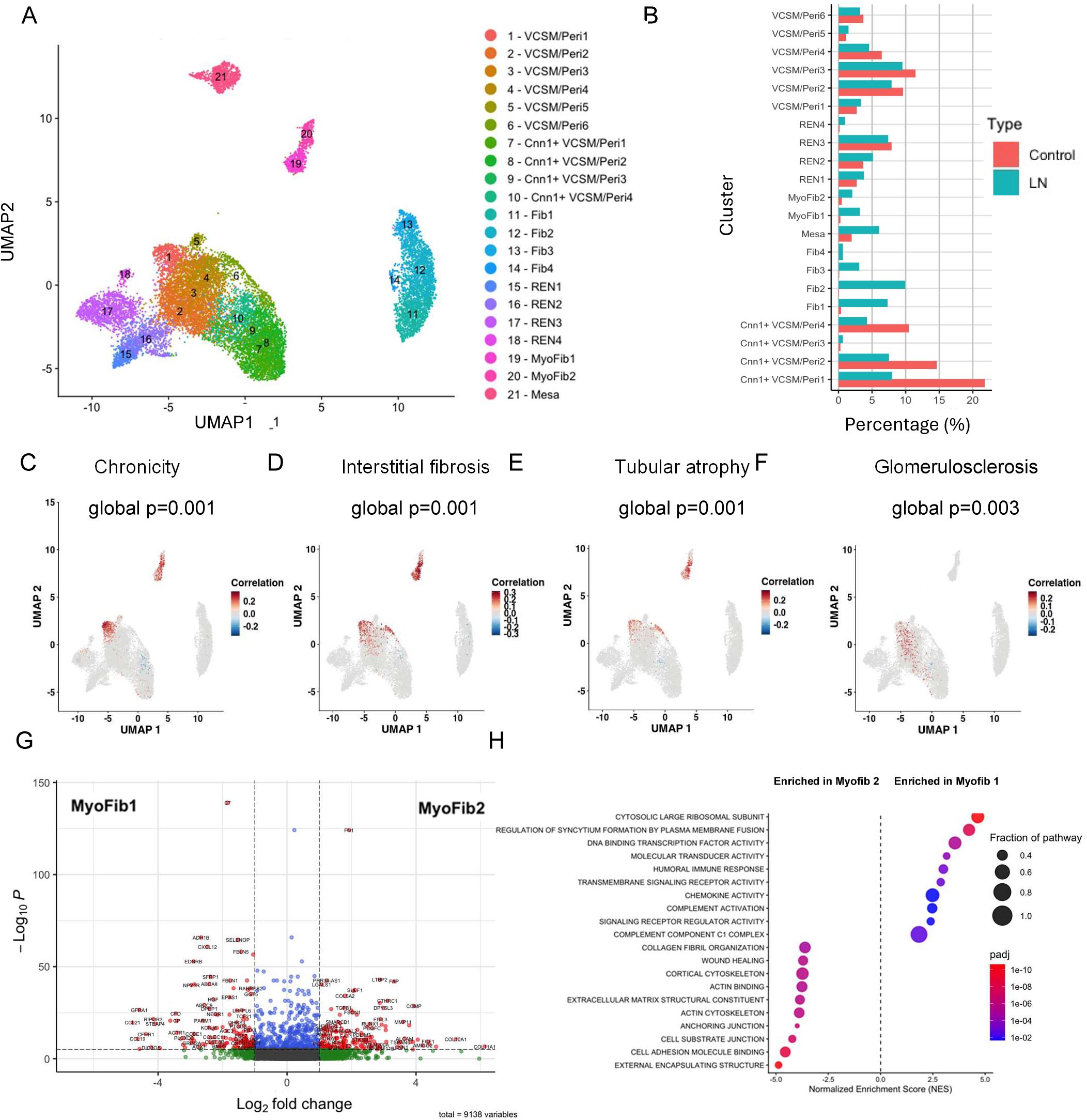
Association of myofibroblasts with disease chronicity scores in LN. **A.** Fine subclustering of stromal cells from 156 LN and 30 healthy donor control kidney biopsies. **B**. Increased frequency of fibroblasts and myofibroblasts among stromal cells in LN compared with biopsies from live donors. **C-F**. CNA analysis shows a significant association of myofibroblasts and CNN VCSM/Peri1 with global chronicity (C) and interstitial fibrosis/tubular atrophy subscores (D, E). **F**. Glomerulosclerosis subscore of chronicity is significantly associated with mesangial cell loss. **G, H.** Volcano plot showing differential gene expression of Myofib1 vs. Myofib2 clusters (G) and corresponding pathway analysis (H).

Myofibroblasts were distinguished from fibroblasts by expression of myosin genes and Acta2 (**Supplementary Figure 2I, Supplementary Table 3**) and comprised two subclusters **(Figure 1A, Supplementary Table 4**). Myofib1 exhibited a pro-inflammatory profile characterized by increased expression of complement genes C1r, C1s and C7, the inflammatory chemokine CXCL12, and ADH1B, an enzyme associated with inflammatory cancer-associated fibroblasts ^16^. Conversely, Myofib2 expressed FAP, COMP, TIMP1, MMP11 and collagens 1A1, 1A2 and 3A1, consistent with a fibrotic/remodeling profile, confirmed by pathway analysis (**Figure 1G, 1H, Supplementary Table 5**).

We next examined the gene expression profile of the Cnn1^-^ VSMC/Pericyte1 cluster that correlated with the chronicity index (**Figure 1D**) versus the other 5 Cnn^-^ VSMC/pericyte clusters (**Supplementary Figure 2J, Supplementary Table 6**). Differentially upregulated genes in the Cnn^-^ VSMC/Pericyte1 cluster include complement genes, collagen genes and inflammatory chemokines as well as STEAP4, a marker for de-differentiated VSMCs ^17^, endothelin receptor B, angiotensin receptor 1, PDGFRB, ITGA1, an adhesion molecule induced by TGFβ, CD36, that is upregulated in pericytes during AKI to CKD progression ^18^, and the osteopontin (Spp1) ligand CD44 whose expression in pericytes is associated with microvascular degeneration ^19^. Pathway analysis confirmed these findings and further showed a decrease in contractile pathways (**Supplementary Table 7**) including loss of the pericyte marker Myh11. These findings suggest that Cnn^-^ VSMC/Pericyte1 represents dedifferentiated microvascular pericytes.

Fine mapping of renal myeloid cells from the AMP/SLE cohort revealed 24 subclusters (**Supplementary Figure 3A**) ^13^. M8, M10, M16, M17 and M18 correspond to C2 macrophages, clusters M0 and M9 contain RM1, and clusters M0, M5 and M9 contain RM0. Clusters M7 and M10 overlap both C2 and RM0 (**Supplementary Figure 3B-E**), supporting the hypothesis of convergent differentiation of the macrophage subsets expressing a damage profile ^13^. Clusters M5, M7, M8, M10, M6, M17 and M18 were significantly enriched in LN biopsies, and we refer to them as disease-associated macrophages ^13^.

To identify the spatial relationships of these macrophage subsets to fibroblasts we analyzed Xenium spatial transcriptomic data in kidney biopsies from 6 LN patients and 2 healthy donors ^12, 13^. We first identified the stromal cells in the spatial transcriptomic dataset by mapping to a reference data set ^2^, then used the AMP reference data set to identify stromal subsets by coarse (**Supplementary Figure 4A**), followed by fine subclustering (**Figure 2A**).

**Figure 2.**
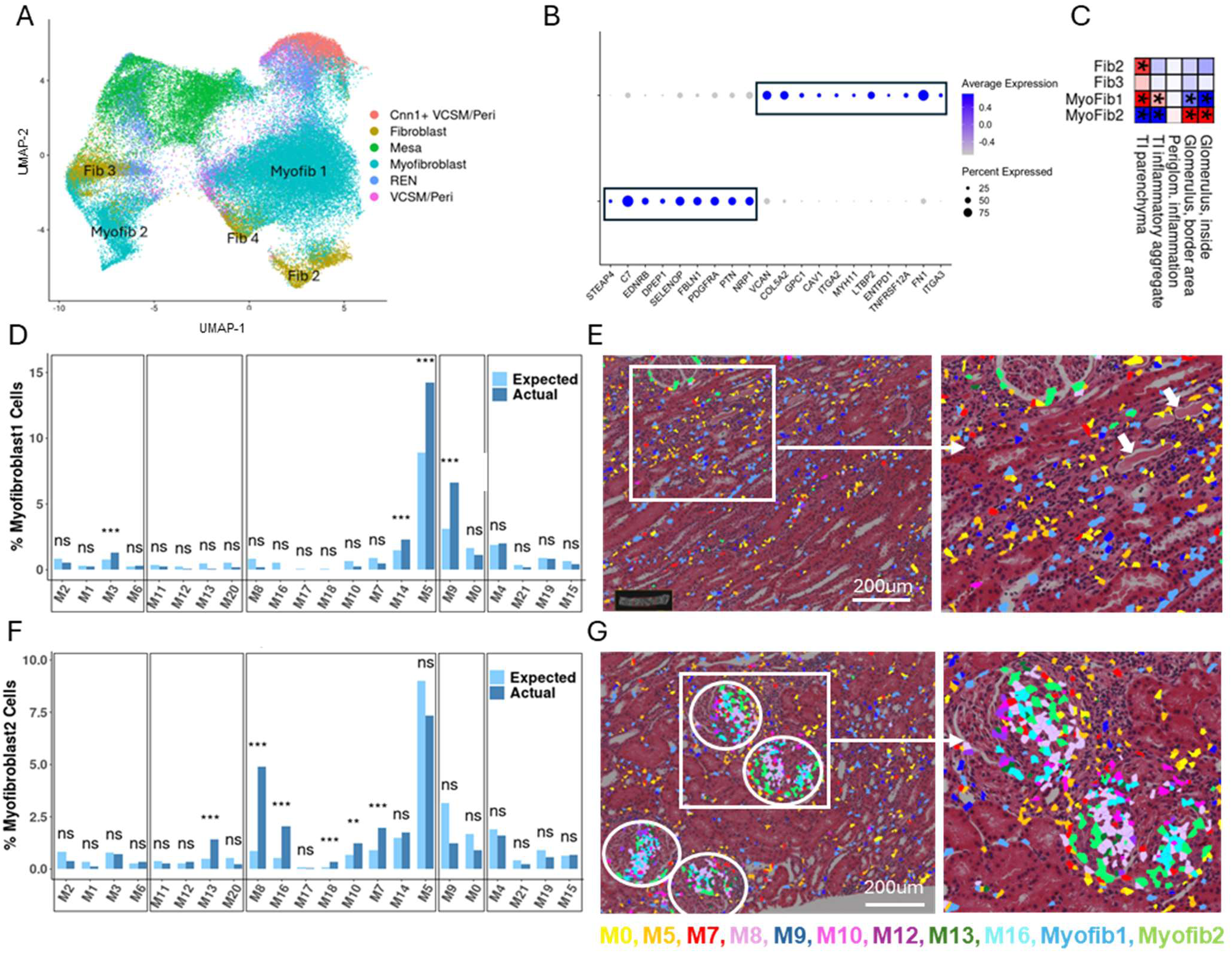
Spatial distribution of myofibroblast subsets in LN biopsies. **A.** Mapping of stromal subclusters. **B.** Spatial transcriptomics markers that distinguish Myofib1 and Myofib2. **C.** The statistical significance of localization preferences, in the 6 analyzed LN kidneys. Each column represents a specific region; each row represents a cluster. Heatmap elements are colored based on the magnitude of the adjusted p values, and the direction of observed deviance from the null distribution assuming no preference: red represents a relative preference of a cluster to localize in a specific region type; blue represents a relative avoidance. Asterisks represent cases with FDR ≤ 0.05, further supported by the individual analysis of at least 2 separate study subjects. **D.** Adjacency of Myofib1 at 10uM distance to macrophage subsets in kidneys of 6 analyzed LN patients. Bars represent cell subsets and show the probability of proximity to each subset. Clusters M1, 2, 3 and 6 are monocytes, Clusters M11,12,13 and 20 are differentiated monocytes, Clusters M5, 7, 8, 10,14, 16, 17 and 18 are macrophages that are present only in LN biopsies, Clusters M0, and 9 are resident macrophages and Clusters M4,15,19 and 21 are dendritic cells ^13^. **E.** Representative LN biopsy showing distribution of Myofib1 (blue) and Myofib2 (green) and their relation to interstitial myeloid cell subsets. White arrows indicate tubules containing casts. **F.** Percent of Myofib2 within 10uM of the indicated macrophage subsets in kidneys of 6 analyzed LN patients. **G**. Representative LN biopsy showing distribution of Myofib2 (green) and their relation to glomerular myeloid cell subsets. Circles indicate glomeruli.

Myofibroblasts were subclustered separately (**Figure 2B, Supplementary Figure 4B, Supplementary Table 4**) and the two myofibroblast subsets were mapped onto the spatial data as previously described ^13^ together with the 24 macrophage subclusters.

Adjacency analysis ^13^ showed that Myofib1 were enriched in the tubulointerstitium, suggesting that they arise from interstitial fibroblasts or pericytes, whereas Myofib2 localized to glomeruli and glomerular borders (**Figure 2C**), suggesting that they may be derived from glomerular stromal cells that include mesangial cells, or from periglomerular stromal cells.

Consistent with these data, Myofib1 colocalized with RM subsets M5 and M9 and, to a lesser extent, M3 and M14 in the interstitium and around lymphocytic infiltrates. (**Figure 2D, E, Supplementary Figure 4C**). By contrast, Myofib2 colocalized with multiple subsets of glomerular infiltrating disease-associated macrophages (**Figure 2F, G, Supplementary Figure 4D**).

To further validate these data, we stained 6 human LN kidneys with antibodies to FAP, expressed by Myofib2 (**Figure 2B**), Mertk, expressed by Cluster M9 and disease-associated macrophages, Trem2, expressed by Cluster M0 and disease-associated macrophages and GPNMB, expressed mainly by disease-associated macrophages (**Figure 3A-D**). FAP expression was observed in 4 of the 6 biopsies and localized to the glomeruli and glomerular borders in Acta2^-^ cells adjacent to macrophages expressing Mertk, Trem2 and/or Gpnmb (**Figure 3E-J**). By contrast, Acta2^+^/FAP^-^ Myofib1 were located in the interstitium in niches containing Mertk+ and - macrophages and scattered lymphocytes (**Figure 3K**).

**Figure 3.**
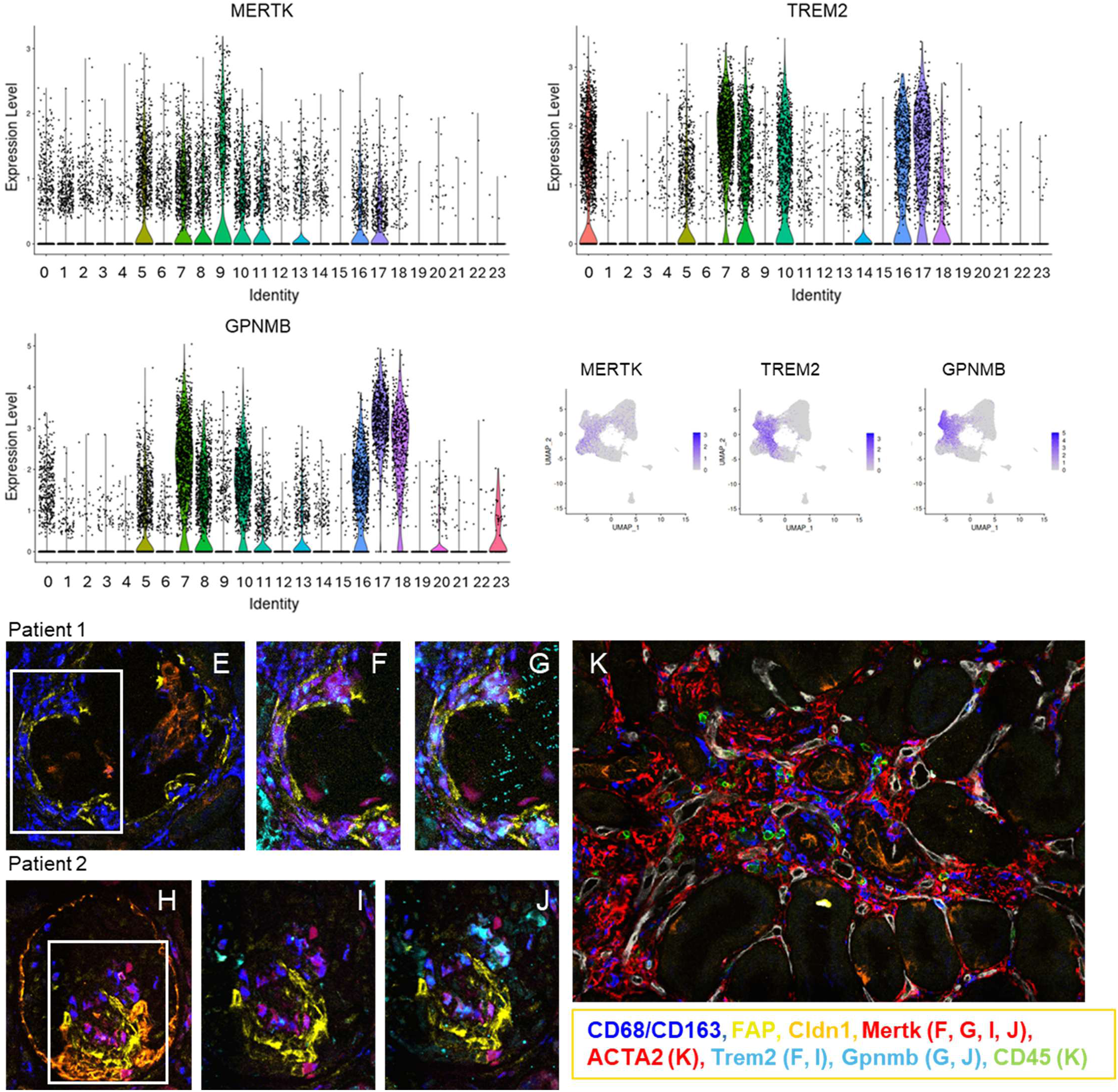
**Immunohistochemistry validation of spatial transcriptomic analyses. A-C**. Violin plots show expression of Mertk (A), Trem2 (B) and Gpnmb (C) in macrophage clusters. **D.** Distribution of cells expressing Mertk, Trem2 and Gpnmb in the macrophage clusters. **E-J.** Glomeruli from 2 representative patients stained with the indicated antibodies show FAP+ myofibroblasts adjacent to Mertk/Trem2+and Mertk+/Gpnmb+ macrophages and areas of PEC proliferation. K. Interstitial fibrogenic niche shows FAP-ACTA2+ myofibroblasts adjacent to Mertk-/Trem2- macrophages and lymphoid cells surrounding injured tubules expressing Cldn1.

Given the difficulty in isolating human LN macrophages for functional studies, we used a mouse coculture system to investigate whether RMs and infiltrating C2 macrophages induced different myofibroblast profiles or functions. We first validated the phenotype of C2 and RMs using flow cytometry for canonical markers (**Supplementary Figure 5A)**. The percentage of RMs positive for both Trem2 and CD9 was higher in nephritic than in prenephritic mice of both strains as was the MFI of Trem2 (**Supplementary Figure 6A, B**), with no change in MFI of either CD9 or CD63 (**Supplementary Figure 6C, D**). C2 cells were expanded in nephritic mice of both strains **(Supplementary Figure 6E**). The MFI of both Trem2 and CD9 was higher in C2 (Ly6C^int^/Ccr2^lo^/Dectin^hi^) than in C1 (Ly6C^hi^/Ccr2^hi^) cells regardless of disease stage (**Supplementary Figure 6F, G**) with no difference in the MFI of CD63 (**Supplementary Figure 6H**).

Freshly isolated CD11b^+^/Ly6G^-^/Ly6C^-^/F4/80^+^/CD81^hi^ RMs from prenephritic and nephritic NZB/WF1 kidneys were cocultured in direct contact with GFP^+^ NZB/WF1 embryonic fibroblasts (MEFs) (**Figure 4A**). RMs from nephritic but not from prenephritic mice induced expression of the fibroblast activation marker FAP on Thy1^+^ fibroblasts (**Figure 4B, Supplementary Figure 6B**). To evaluate the role of secreted factors by both cell types in the cocultures, we collected 48hr conditioned media (CM) from the cocultures and stimulated fresh MEFs for an additional 48hrs (**Figure 4A**). FAP induction was lower than that observed in direct cocultures nephritic CM, but with a trend towards higher MFI in nephritic compared with prenephritic CM-treated MEFs (**Figure 4C**).

**Figure 4.**
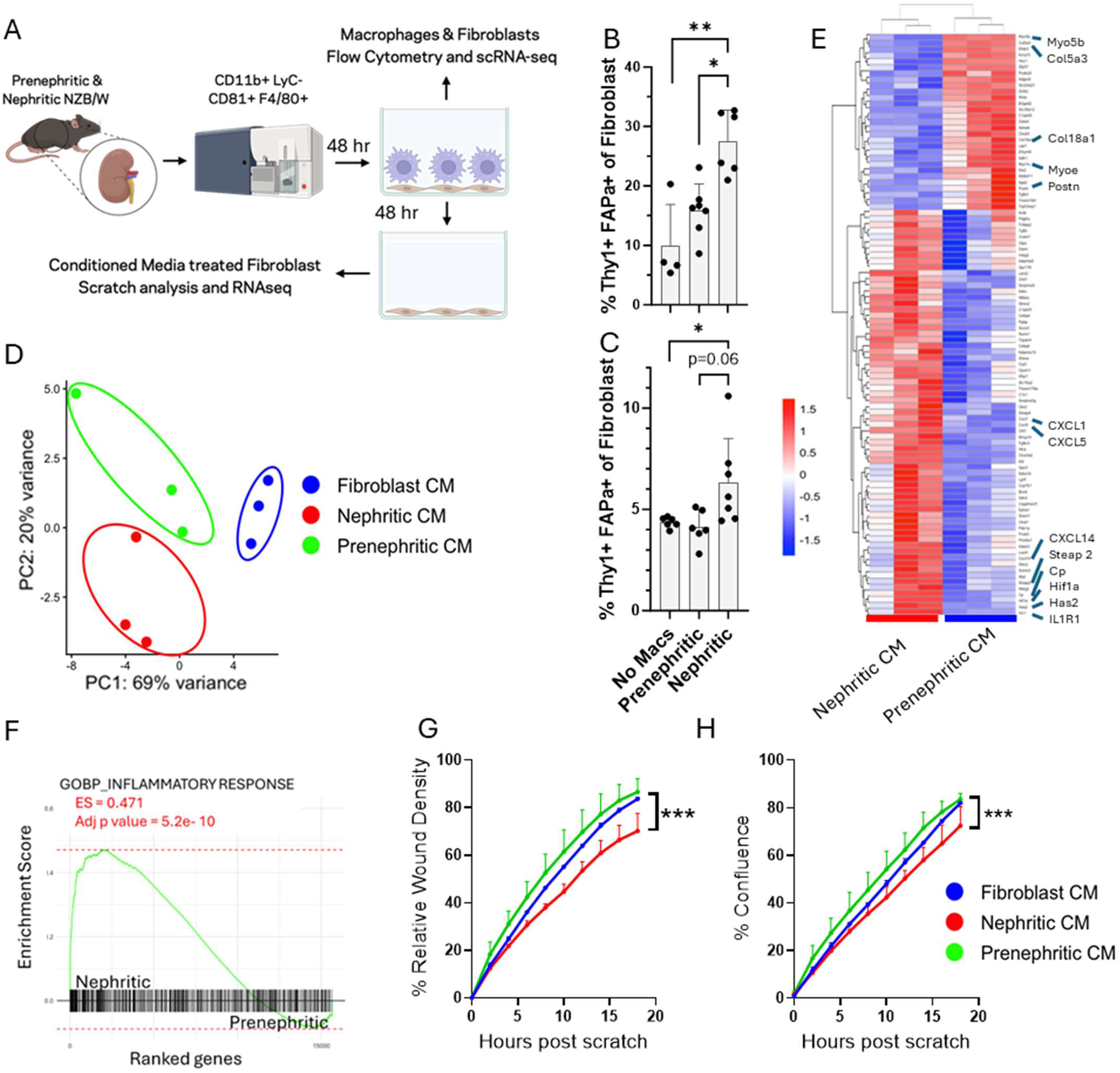
Coculture of resident renal macrophages with mouse embryonic fibroblasts. **A.** Experimental scheme using direct cocultures and cocultures with conditioned media (CM). **B, C**. Induction of FAP on Thy1+ fibroblasts after coculture with prenephritic or nephritic resident renal macrophages (B) or conditioned media (CM) from 48-hr cocultures (C). Each symbol indicates an individual mouse. Kruskal-Wallis ANOVA with Dunn’s correction for multiple comparisons *p<0.05: **p<0.01. **D.** PCA analysis of fibroblasts exposed to CM from nephritic or prenephritic resident macrophages or fibroblasts alone. **E.** Heat map showing differences in gene expression between MEFs cocultured with nephritic or prenephritic resident macrophage conditioned media. **F.** GSEA analysis indicates inflammatory response as a major pathway differentiating the MEFs exposed to nephritic compared with prenephritic CM. **G, H.** Conditioned media from resident macrophage/MEF cocultures decreases wound density (G) and wound confluence (H) in a scratch assay. Data represent the mean +/- SD of 3 mice per group One way ANOVA with Friedman’s test, ***p<0.001.

PCA analysis of bulk RNAseq showed clear separation of MEFs exposed to CM from prenephritic compared with nephritic RMs and both differed from MEFs exposed to CM from fibroblasts alone (**Figure 4D**). CM from prenephritic RM cocultures induced a contractile profile including collagen genes, Myo5b, Myo1b, Hes1 and Postn. By contrast, CM from nephritic RM cocultures induced proinflammatory chemokines CXCL1 and CXCL5, IL1R, the mesenchymal marker Steap2 and its interacting protein Cp ^20^ (**Figure 4E, Supplementary Table 8**) and the fibrotic regulators Hif1a and Has2 ^21, 22^. This proinflammatory gene expression pattern was confirmed by pathway analysis (**Figure 4F, Supplementary Table 9**).

To explore the functional consequences of these differences, we subjected MEFs to a scratch wound and added 48hr CM from the cocultures (**Figure 4A**). CM from prenephritic RM cocultures and from MEFs alone induced similar wound healing kinetics and wound confluency but these parameters were impaired in the MEFs exposed to CM from nephritic RMs (**Figure 4G, H**). This observation is consistent with a remodeling role of inflammatory fibroblasts ^6^ and our previous data showing an increase in the *in vivo* activity of degradative enzymes (MMPs and Cathepsins) in nephritic compared with prenephritic RMs from NZB/W mice ^23^.

To further understand how direct contact with RMs influences fibroblast differentiation, cells from the direct cocultures (**Figure 4A**), were harvested for scRNAseq. We identified 2 major subclusters of macrophages distinguished by expression of Spp1 (**Figure 5A-F**).

**Figure 5:**
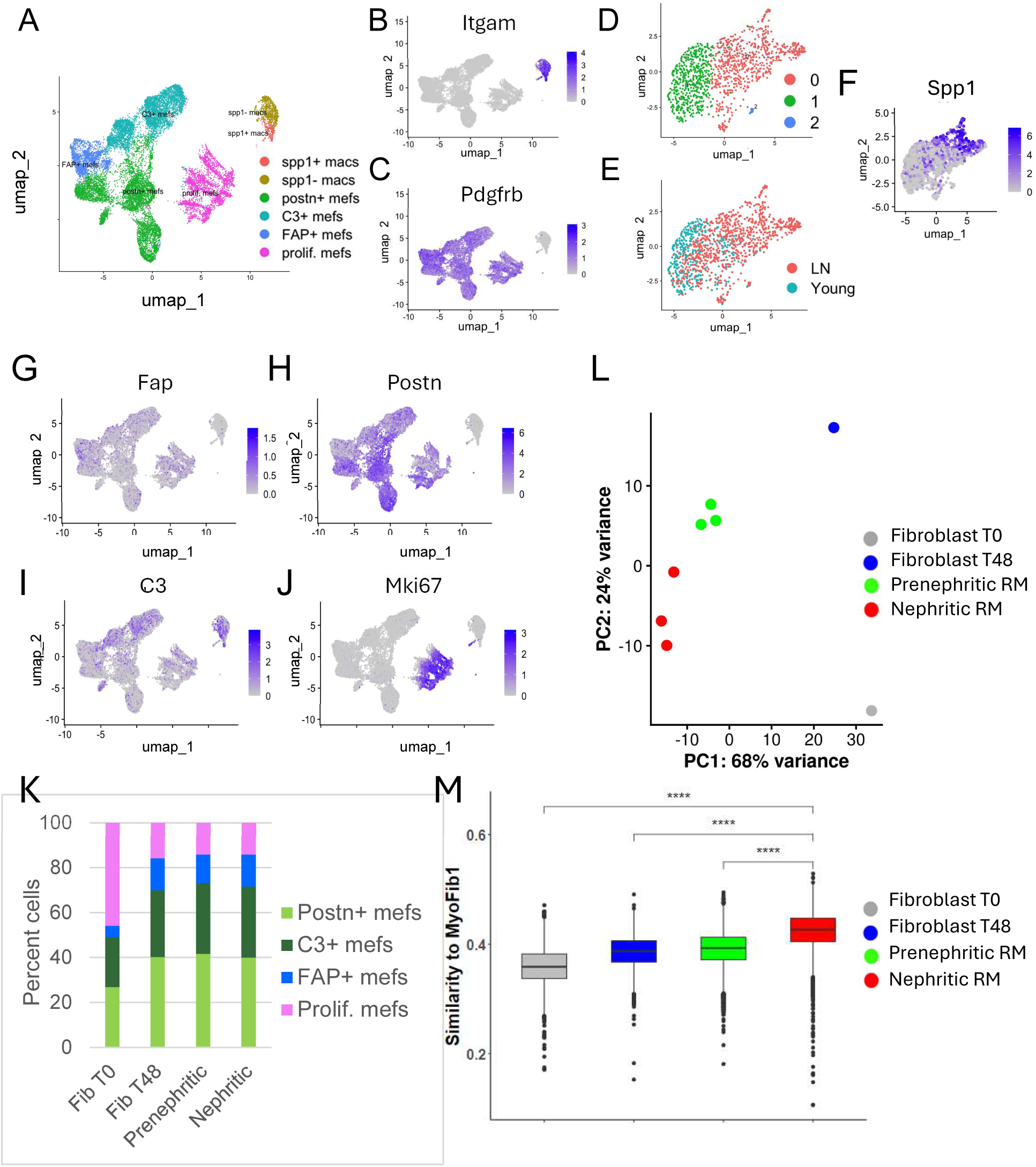
**Subcluster analysis of single cells from cocultures of resident macrophages and MEFs at 0 and 48 hours**. **A-C.** UMAP of cocultured cells shows 2 subclusters of macrophages and 4 subclusters of fibroblasts distinguished by expression of CD11b (B) and PDGFRB (C). **D-F.** Fine clustering of macrophages shows 2 major clusters differentiated by disease stage (E) and expression of Spp1 (F). **G-J.** Feature maps show fibroblast subclusters differentiated by expression of Fap, Postn, C3 and Ki67. **K.** No difference in the distribution of cells in each differentiated fibroblast cluster between the 3 groups of 48hr cocultures. By contrast, Time 0 fibroblasts have a prominent proliferating cluster. **L.** Pseudobulk analysis of scRNAseq data. Cells from T0 and T48 samples (3 per group) were pooled for scRNAseq analysis and cells from 3 prenephritic and nephritic mouse co-cultures were analyzed individually. **M.** MEFs cocultured with nephritic RMs are more similar to Myofib1 than MEFs cocultured with prenephritic RMs or MEFs cultured alone. **** p<0.0001.

Differential gene expression analysis confirmed that Spp1^+^ macrophages shared genes that distinguish nephritic from prenephritic RMs by bulk RNAseq (**Supplementary Table 10**) indicating that they retained their disease-associated profile in culture.

We classified the cocultured MEFs into four major subclusters exemplifying the heterogeneity of possible differentiation pathways. Time 0 MEFs were predominantly found within the proliferating cluster. By 48hrs, however, MEFs exposed to RMs, or cultured alone, differentiated into C3^+^, Fap^+^, and Postn^+^ subsets (**Figure 5G-J**,) whose cluster frequency was proportionally similar in all 3 groups of 48hr cocultures, (**Figure 5K**). Differential gene expression and GSEA analysis (**Supplementary Tables 11 and 12**) revealed that C3^+^ MEFs exhibited an ECM signature. Fap^+^ MEFs were characterized by expression of genes associated with collagen fibril organization and a distinct set of transcription factors such as Twist1 and Foxd1, both previously associated with profibrotic function ^24, 25^. Postn^+^ MEFs expressed genes linked to myofibril assembly and contraction, as well as autophagy, a process reported to suppress myofibroblast differentiation ^26^. All three subsets showed downregulation of genes involved in cell division.

Differential gene expression analysis and pseudobulk analysis (**Figure 5L**) revealed that MEFs cocultured with RMs from nephritic mice upregulated genes associated with inflammatory cytokine and chemokine signaling, ECM organization and collagen biosynthesis while downregulating genes associated with cell cycle (**Supplementary Tables 13, 14**). All subclusters of MEFs cocultured with RMs from nephritic mice were more similar to Myofib1 than those from the other coculture conditions (**Figure 5M**).

To further explore the interactions between MEFs and RMs we performed CellChat analysis. All 3 differentiated MEF clusters interacted with macrophages via Csf1 (**Supplementary Figure 7A**). Spp1^+^ macrophages preferentially interacted with MEFs through the IL1 and Sema4 pathways (**Supplementary Figure 7B, C**) whereas Spp1^-^ macrophages interacted with MEFs via the Osm pathway (**Supplementary Figure 7D**). Both macrophage subsets communicated with MEFs via the PDGF and TNF pathways (**Supplementary Figure 7E, F**).

To validate these interactions in the human LN biopsies, we performed CellChat analysis of interactions between the two human myofibroblast subsets and their adjacent macrophage clusters (**Figure 2D, F**). We identified interactions between Myofib1 and RM cluster M5 via Spp1/integrins, tubulointerstitial macrophage clusters 3 and 14 via Sema4/PlexinB2, and RM cluster M9 via PDGF/PDGFR (**Supplementary Figure 7G-I, Supplementary Figure 8A**).

CellChat analysis of interactions between Myofib2 and their adjacent glomerular macrophage clusters (**Figure 2F**) confirmed that Spp1/integrin αVβ1 interactions were dominant (**Supplementary Figure 7G**). Additional pathways included the CD49d (Itgα4/Itgβ1)/VCAM and Visfatin (NAMPT - nicotinamide phosphoribosyl transferase)/INSR, pathways for Myofib1 and the Fn1, Granulysin/Sort1 and CD99/CD99 pathways for Myofib2 (**Supplementary Figure 7J-M, Supplementary Figure 8B**).

We were unable to isolate sufficient primary C2 macrophages from nephritic mice for coculture assays due to their low frequency compared with RMs. A damage-associated macrophage phenotype can be induced by exposing bone marrow derived macrophages (BMDM) to a variety of *in vitro* conditions including GM-CSF + IL17, and even to a certain extent by culture with GM-CSF alone ^27^. GM-CSF + IL17 induced higher expression of CD9 and Trem2 compared with either classical IFNγ/LPS M1 or IL4/IL13 M2 polarization, (**Figure 6A, B**). To further validate the C2-like phenotype of IL-17 polarized BMDMs we constructed a similarity score for C2 renal macrophages using bulk RNAseq data from the polarized macrophages and the previously reported scRNAseq gene C2 expression profile ^12^. This analysis confirmed that IL17 polarized BMDMs had the highest similarity to C2 macrophages (**Figure 6C**).

**Figure 6.**
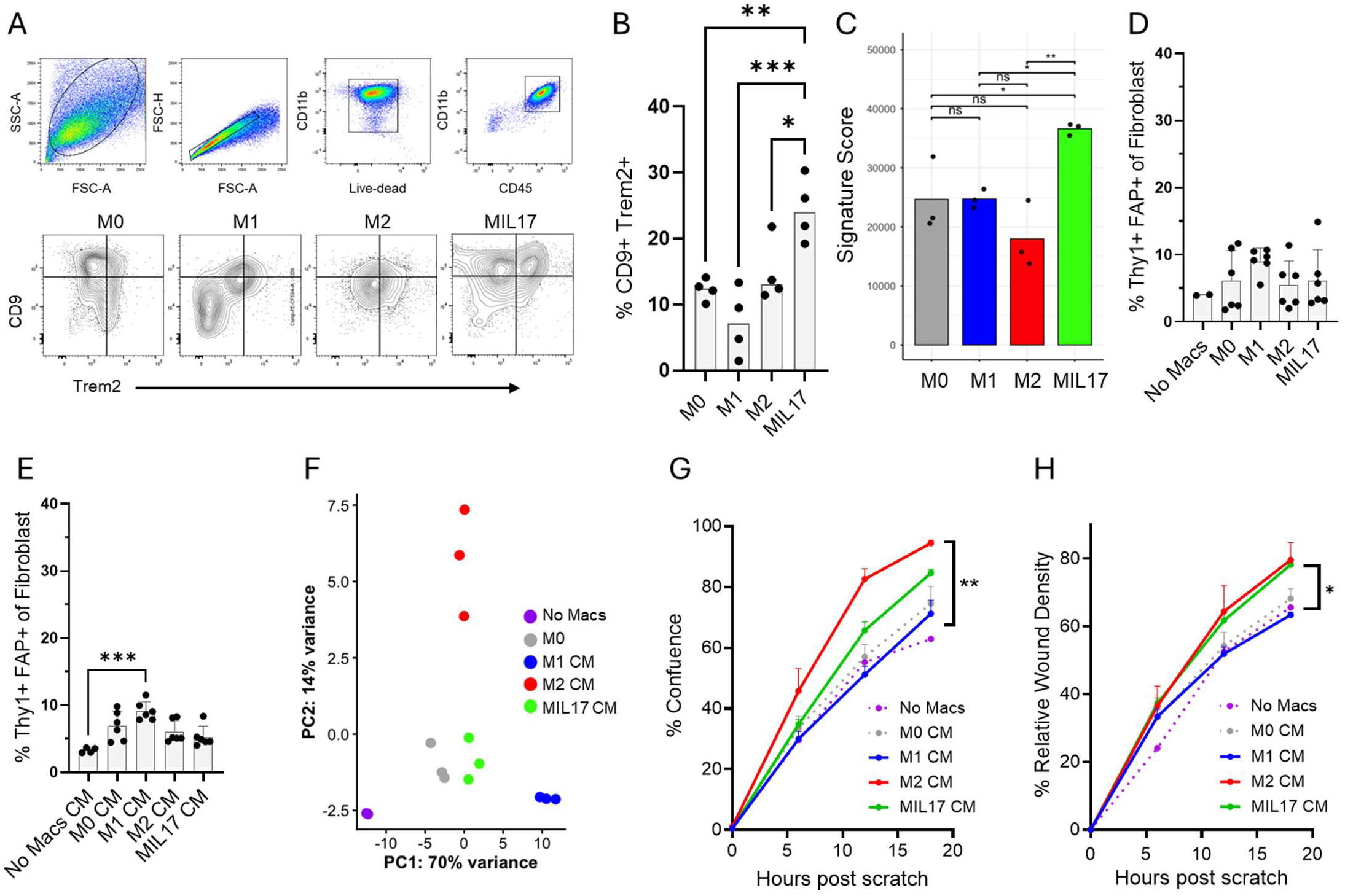
**Induction of a damage-associated macrophage phenotype with GM-CSF and IL17**. **A, B**. Flow cytometry of BMDM exposed to GM-CSF and polarizing cytokines shows induction of a CD9+/Trem2+ phenotype by IL17. **C.** Similarity score of polarized macrophages to renal C2 macrophages. **D, E.** Induction of FAP on Thy1+ fibroblasts after direct coculture with polarized macrophages (D) or conditioned media (CM) from 48hr cocultures (E). **B, D, E.** Each symbol indicates an individual mouse. Kruskal-Wallis ANOVA with Dunn’s correction for multiple comparisons *p<0.05, **p<0.01, ***p<0.001. **F.** PCA analysis of genes from MEFs exposed to polarized macrophages. **G, H.** Conditioned media from MIL17 polarized macrophage/MEF cocultures increased wound confluence (F) and wound density (G) in a scratch assay. Data represent the mean +/- SD of 3 mice per group. One way ANOVA with Friedman’s test *p<0.05, **p<0.01.

Bulk RNAseq profiling confirmed the expected transcriptional profiles of polarized M1 and M2 macrophages relative to M0 macrophages (**Supplementary Table 15**). Both M0 and MIL17 macrophages expressed higher levels of Trem2, Spp1, Fabp5, and Gpnmb than M1 or M2 macrophages. However, compared with M0 macrophages, IL17 exposure induced upregulation of genes associated with the acute phase response, inflammation, reactive oxygen species production, ECM organization, and cell chemotaxis, including CXCL1, CXCL5, Nos2, IL6, Saa3, and Yap1 (**Supplementary Tables 15 and 16**). Pathway analysis supported these findings (**Supplementary Table 17**).

Minimal FAP induction was induced on MEFs that were cocultured with polarized BMDMs or their CM with no difference among the macrophage conditions (**Figure 6D, 6E**). Nevertheless, bulk RNAseq profiling revealed significant transcriptional differences between MEFs exposed to the CM from polarized macrophages (**Figure 5F, Supplementary Tables 18-20**). GSEA analysis showed that compared to M0 CM, both M1 and MIL17 CM induced an inflammatory fibroblast pattern and downregulated cell cycle genes. M1-induced fibroblasts exhibited a stronger inflammatory gene signature than MIL17-induced fibroblasts, which instead showed enrichment for genes involved in cytoplasmic translation and organ morphogenesis (**Supplementary Tables 18, 19**). Compared to M0, M2-induced fibroblasts expressed genes associated with RNA processing and cytoplasmic translation and downregulated genes associated with extracellular structure. MIL17-induced fibroblasts were less proliferative than M2-induced fibroblasts but expressed more genes associated with extracellular structure (**Supplementary Tables 18, 20**).

To evaluate the functional effects of the polarized macrophages on fibroblasts we exposed MEFs to a scratch wound in the presence of CM. Wound confluence was enhanced by M2 CM but not by M1 CM, while the MIL17 CM had an intermediate effect (**Figure 6G**). By contrast, while M1 CM did not enhance wound density, both M2 and MIL17 CM enhanced wound density to the same degree (**Figure 6H**). These data show that MIL17-induced fibroblasts promoted wound healing but were less effective at wound closure than M2-induced fibroblasts, perhaps due to their lower proliferative capacity.

## DISCUSSION

Progression of LN from active to chronic disease is associated with poor response to treatment, declining kidney function, and renal fibrosis. Myofibroblasts are the primary ECM-producing fibroblast subset responsible for fibrotic tissue remodeling and stiffening ^10^. We show here that their frequency in LN renal biopsies correlates with the histologic chronicity score, highlighting the need to better understand how myofibroblasts are induced in LN kidneys.

Among disease-associated fibroblast subsets, inflammatory fibroblasts can attract immune cells and promote formation of tertiary lymphoid structures in inflammatory tissues. By contrast, fibroblasts expressing the serine protease FAP (fibroblast activation protein) are associated with collagen cleavage, ECM remodeling, tissue stiffening and immunosuppressive functions ^28^, promoting immune cell exclusion. Alternatively, remodeling fibroblasts produce excess ECM-degrading enzymes that destroy surrounding tissues ^6, 7^. A dichotomy of fibroblast states can exist in inflammatory tissues in which spatially separate inflammatory CCL19^+^CXCL10^+^ fibroblasts and remodeling SPARC^+^Col3A1^+^ fibroblasts both associate with inflammation ^6^. Although it has been suggested that these two broad subsets are developmentally related, delineating their origins in the kidney will require further investigation.

Pericytes are a major source of myofibroblasts during inflammation ^29^ and can secrete inflammatory cytokines, contributing to fibrosis. We show here that alterations in the frequency of pericyte subsets correlate with chronicity in human LN, including the association of mesangial cell loss with glomerulosclerosis and increased Cnn^-^ VSMC/pericyte1 frequency with chronicity. Differentially expressed genes in Cnn^-^ VSMC/pericyte1 from LN kidneys are associated with pericyte dedifferentiation and detachment, as well as induction of chemokines that could recruit inflammatory cells.

Macrophages are present in the renal fibrogenic niche ^2, 30^ and play a critical role in tissue repair, inflammation, and ECM remodeling. Following injury, macrophages help clear cellular debris, promote angiogenesis, and stimulate fibroblast activation and ECM remodeling to restore tissue integrity. However, in chronic conditions, macrophages can drive fibrosis through sustained fibroblast activation, leading their differentiation into myofibroblasts. Additionally, inefficient macrophage efferocytosis occurs in LN ^12^, perpetuating inflammation. The balance between reparative and fibrotic/inflammatory macrophage activity dictates whether healing proceeds effectively or results in fibrosis.

Our previous work using LN kidney biopsies identified a cluster of infiltrating glomerular macrophages (Classical 2), found in proliferative LN, that correlates with histologic disease activity but declines with chronicity ^12^. Classical 2 macrophages comprise several sub-clusters ^13^ that variably express genes characteristic of a specialized subset of macrophages that accumulates in the fibrotic niche across organs in response to chronic inflammation, tissue damage, and metabolic stress ^11, 14, 31^. Their functional role has been a subject of debate, since deletion of several of their canonical genes such as Trem2 and Gpnmb exacerbate organ injury in fibrosis models ^32, 33^. By contrast, deletion of the canonical gene Spp1 (osteopontin) appears to protect from fibrosis in multiple animal models, including in the kidney ^34^. Recent studies suggest that a differentiated subset of these cells, marked by higher Spp1 expression and the acquisition of inflammatory chemokines, confers pro-fibrotic function and pathogenicity ^35^.

Spatial transcriptomic analysis revealed adjacency of kidney-infiltrating subclusters of these disease-associated macrophages to glomerular myofibroblasts (Myofib2) located inside glomeruli and at the glomerular borders within a microenvironment that also included expanded layers of parietal epithelial cells ^13^. Macrophages with a similar gene expression pattern were generated *in vitro* by exposure to GM-CSF and IL17 ^36^, enabling coculture studies with undifferentiated fibroblasts. MIL17-cocultured fibroblasts exhibited dysfunctional wound healing, characterized by equivalent wound density but incomplete closure compared to reparative M2-cocultured fibroblasts. These findings are consistent with their enhanced expression of fibrotic and ECM genes together with reduced expression of proliferative genes. The source of Myofib2 in the glomeruli of LN patients is not yet clear as glomerular PDGFRB^+^ cells, that include the mesangial cell population, are heterogeneous ^37^. We hypothesize that as mesangial cells decrease and Myofib2 emerge with chronicity, the ensuing structural injury to capillaries together with increasing scar tissue may limit immune cell infiltration or survival, driving the observed decline in infiltrating glomerular macrophages associated with disease chronicity ^12^.

We also previously identified a population of resident renal macrophages (RM0) that expand during both proliferative and membranous LN and correlate with the chronicity score ^12^. These cells acquire an activated gene expression profile partially overlapping with that of infiltrating disease-associated macrophages including upregulation of Spp1 and Trem2. Activated resident macrophages with this profile (Cluster M5) localize mainly to the tubulointerstitium and inflammatory infiltrates where they interact with Myofib1, but they were also observed in periglomerular areas where they may differentiate further ^13^ into M7 and M10 clusters and interact with Myofib2. Importantly, our coculture studies show that resident macrophages from nephritic mice drive an inflammatory and remodeling fibroblast profile. The induction of this remodeling phenotype is also consistent with the high production of MMPs by nephritic resident macrophages ^23^ and the impaired wound closure observed following coculture of fibroblasts with nephritic macrophage CM. Although fibroblasts co-cultured with nephritic RMs are most likely to acquire this Myofib1-like program, we show here, using scRNAseq analysis of direct cocultures, that alternate differentiation programs are possible. More experiments are needed to determine whether, in addition to glomerular stromal cells, activated periglomerular RMs help to drive FAP production at the glomerular borders.

A cardinal feature of disease-associated macrophages is their acquisition of Spp1 expression. Spp1 encodes the ECM protein osteopontin that is required for myofibroblast differentiation ^34, 38^. It interacts with surface adhesion molecules on fibroblasts including αv integrins and CD44 and enhances TGFβ signaling while preventing fibroblast apoptosis. We identified prominent Spp1/adhesion molecule interactions between macrophages and both Myofib1 and Myofib2 in LN patients, indicating that Spp1 mediates disease-associated macrophage/myofibroblast interactions across both glomerular and interstitial compartments.

By employing coculture studies and CellChat analyses, we found that in addition to Spp1, IL1 and Sema4A/D are implicated in macrophage-fibroblast interactions. Myeloid cells are a major source of IL1 in LN kidneys ^39^ and IL1 is known to delay wound closure in other models ^40^. Monocyte Sema4A production is reported in scleroderma and can induce IL17 production by T cells ^41^. Both Sema 4A and Sema 4D induce inflammatory mediator secretion and ECM deposition by fibroblasts through direct engagement of Plexins including Plexin B2 ^41, 42^. By contrast, Osm (Oncostatin M), a key mediator of interactions between fibroblasts and Spp1^-^macrophages, promotes a regenerative ECM remodeling program in injury models ^43^. In the LN biopsies NAMPT, an inflammatory adipocytokine that promotes fibrosis and fibroblast senescence ^44^, and VCAM, an adhesion molecule made by inflammatory cancer-associated fibroblasts that enhances cellular invasion ^45^, mediate interactions between interstitial macrophages and Myofib1. By contrast, CD99, an adhesion molecule that mediates macrophage transmigration ^46^ and granulysin that engages the profibrotic membrane protein sortilin mediate interactions with Myofib2 ^47^. Notably, antagonism of several of these pathways attenuates fibrosis in animal models ^46, 47, 48^.

Despite the insights gained, there are several limitations of this study. *In vitro* coculture systems, while providing mechanistic insight, may not fully recapitulate the complexity of the *in vivo* renal microenvironment, including the influence of other stromal interactions, mechanical forces and 3D structure. By isolating fibroblasts as the sole stromal component, we may overlook effects from other stromal cells that shape macrophage behavior or respond differently to immune-derived signals. Furthermore, while our data implicate various receptor ligand pairs in macrophage fibroblast interactions, definitive validation using targeted inhibition or genetic models is now required to demonstrate causality. Mouse models, including the two strains used here, recapitulate many aspects of human LN but cannot fully capture its complexity, heterogeneity, and chronicity. The human spatial transcriptomic datasets provided spatial and transcriptomic validation but were limited in patient number, potentially constraining generalizability. Finally, because kidney fibroblasts manifested poor growth characteristics, we used embryonic fibroblasts that share many transcriptional features with kidney fibroblasts but may not behave identically in coculture.

Together, our data reveal that spatially and functionally distinct renal macrophage subsets in LN kidneys drive maladaptive fibroblast activation, linking macrophage–fibroblast crosstalk to both glomerulosclerosis and interstitial fibrosis. Our findings may also have clinical implications. Recent biomarker studies of urine from the AMP-SLE cohort defined a proteomic macrophage signature whose early resolution predicts clinical response ^49^ and further identified persistence of urinary myofibroblast derived Tenascin-C as a significant biomarker of poor long-term outcome ^50^. Urinary Tenascin-C is an established biomarker of CKD progression, and kidney fibrosis across multiple renal diseases ^3^. These findings underscore the clinical relevance of macrophage–myofibroblast interactions in LN and their ability to be evaluated non-invasively. More broadly, the integration of single-cell sequencing and spatial transcriptomics is refining our understanding of the cellular networks that drive irreversible fibrosis and organ damage, providing a framework for the development of therapies aimed at disrupting pathogenic fibro-inflammatory circuits, limiting maladaptive tissue remodeling, and preserving organ function in lupus and other fibrotic kidney diseases.

## ACKNOWLEDGEMENTS AND AFFILIATIONS

We thank the patients who participated in this study, and the scientists and clinical sites in the Accelerating Medicines Partnerships in RA/ SLE. This work was supported by the Accelerating Medicines Partnership® Rheumatoid Arthritis and Systemic Lupus Erythematosus (AMP® RA/SLE) Network (see below). AMP is a public-private partnership (AbbVie, Arthritis Foundation, Bristol-Myers Squibb Company, Foundation for the National Institutes of Health, GlaxoSmithKline, Janssen Research and Development, LLC, Lupus Foundation of America, Lupus Research Alliance, Merck & Co., Inc., National Institute of Allergy and Infectious Diseases, National Institute of Arthritis and Musculoskeletal and Skin Diseases, Pfizer, Inc., Rheumatology Research Foundation, Sanofi and Takeda Pharmaceuticals International, Inc.) created to develop new ways of identifying and validating promising biological targets for diagnostics and drug development. Accelerating Medicines Partnership and AMP are registered service marks of the US Department of Health and Human Services. Funding was provided through grants from the National Institutes of Health: UH2-AR067676 (AMP RA/SLE), UH2-AR067677 (AMP RA/SLE), UH2-AR067679 (AMP RA/SLE), UH2-AR067681 (AMP RA/SLE), UH2-AR067685 (AMP RA/SLE), UH2-AR067688 (AMP RA/SLE), UH2-AR067689 (AMP RA/SLE), UH2-AR067690 (AMP RA/SLE), UH2-AR067691 (AMP RA/SLE), UH2-AR067694 (AMP RA/SLE), UM2-AR067678 (AMP RA/SLE), RO1-DK131482, U19-AI144306, R01-AI1802222.

## AMP RA/SLE Network: Operation/Scientific

Operations – William Apruzzese, Jennifer Goff, Patrick J. Dunn

SBG – Soumya Raychaudhuri, Fan Zhang, Ilya Korsunsky, Aparna Nathan, Joseph Mears, Kazuyoshi Ishigaki, Qian Xiao, Nghia Millard, Kathryn Weinand, Saori Sakaue

LC/STAMP – PJ Utz, Rong Mao, Bill Robinson, Holden Maecker OMRF – Judith James, Joel Guthridge, Wade DeJager, Susan Macwana **SLE**

Rochester – Jennifer Anolik, Jennifer Barnas

BWH – Michael B. Brenner, James Lederer, Deepak A. Rao Northshore – Betty Diamond, Anne Davidson, Arnon Arazi UCSF – David Wofsy, Maria Dall’Era, Raymond Hsu

NYU – Jill Buyon, Michael Belmont, Peter Izmirly, Robert Clancy, Phillip Carlucci, Kristina Deonaraine

Einstein – Chaim Putterman, Beatrice Goilav Hopkins – Michelle Petri, Andrea Fava, Jessica Li MUSC – Diane L. Kamen

Cincinnati – David Hildeman, Steve Woodle

Broad – Nir Hacohen, Paul Hoover, Thomas Eisenhaure, Michael Peters, Arnon Arazi, Tony Jones, David Lieb

Rockefeller – Thomas Tuschl, Hemant Suryawanshi, Pavel Morozov, Manjunath Kustagi UCLA – Maureen McMahon, Jennifer Grossman

UCSD – Ken Kalunian

Michigan – Matthias Kretzler, Celine Berthier, Jeffery Hodgin, Raji Menon Texas Tech – Fernanda Payan-Schober, Sean Connery

Cedars – Mariko Ishimori, Michael Weisman

## AUTHORSHIP CONTRIBUTION STATEMENT

**Chirag Raparia**: Formal analysis, Conceptualization, Methodology, Investigation, Data curation, Writing – original draft, Writing – review & editing, Visualization. **Paul J. Hoover**: Formal analysis, Conceptualization, Methodology, Investigation, Data curation, Writing – original draft, Writing – review & editing, Visualization. **Junting Ai**: Resources, Data curation. **Sujal Shah**; Data curation, Visualization. **Marcus R. Clark**: Funding acquisition, Supervision. The Accelerating Medicines Partnership (AMP) RA/SLE Network: Resources. **Betty Diamond**: Writing – review & editing, Resources. Anne Davidson: Supervision, Conceptualization, Funding acquisition, Data curation, Writing – original draft, Writing – review & editing, Resources. **Nir Hacohen**: Supervision, Conceptualization, Funding acquisition, Writing – review & editing. **Arnon Arazi**: Supervision, Conceptualization, Formal analysis, Writing – original draft, Writing – review & editing, Visualization.

## DATA STATEMENT

All data will be uploaded to public databases upon manuscript publication. In the meantime, all data will be made available to reviewers upon request.

## Key messages

### What is already known on this topic

`o` Disease-associated macrophages (D-Macs) with a canonical gene expression profile have been identified in lupus nephritis (LN) and across other immune-mediated and fibrotic diseases. Despite extensive characterization of macrophage heterogeneity and disease associations in LN across mice and humans, the spatial interactions of D-Macs and their functions within the stromal microenvironment remain understudied.
• What this study adds

`o` Leveraging the Accelerating Medicines Partnership SLE (AMP-SLE) cohort (156 LN patients and 30 healthy controls), we identified two myofibroblast populations in LN kidneys characterized by distinct inflammatory and fibrogenic transcriptional programs. These populations localize to spatially distinct niches within the kidney during LN and are associated with unique D-Mac subsets and cellular interaction networks.
• How this study might affect research, practice, or policy

`o` A better understanding of the composition of transcriptionally and spatially distinct renal niches during LN enables more precise elucidation of stromal–immune communication networks. This study supports the development of more targeted and effective therapeutic strategies for renal fibrosis in the context of LN.

## Notes

### Competing Interest Statement

The authors have declared no competing interest.

