## Supplementary Material for "Spatially Distinct Macrophage Subsets Drive Myofibroblast Heterogeneity and Maladaptive Fibrosis in Lupus Nephritis"

**LIST OF SUPPLEMENTARY TABLES**

Supplementary Table 1: Demographics of the AMP/SLE Cohort
Supplementary Table 2: Subcluster analysis of stromal cells
Supplementary Table 3: Differential gene expression of Myofibroblasts vs. Fibroblasts
Supplementary Table 4: Differential gene expression of Myofib2 vs. Myofib1
Supplementary Table 5: GSEA of Myofib1 vs. Myofib2 (top pathways)
Supplementary Table 6: Differential gene expression of CNN- VCSM/Peri1 vs. other CNN- VCSM/Peri clusters
Supplementary Table 7: GSEA of CNN- VCSM/Peri1 vs. other CNN- VCSM/Peri clusters (top pathways)
Supplementary Table 8: Differential gene expression of MEFs exposed to CM from nephritic vs. prenephritic resident macrophages.
Supplementary Table 9: Pathway analysis of MEFs exposed to CM from nephritic vs. prenephritic resident macrophages.
Supplementary Table 10: Differential gene expression of Spp1+ and Spp1- macrophages
Supplementary Table 11. Differential gene expression of C3, FAP and Post MEFs
Supplementary Table 12: Significantly regulated pathways in MEFs cocultured with resident macrophages.
Supplementary Table 13: Differential gene expression between MEFs cocultured with nephritic vs. prenephritic resident macrophages.
Supplementary Table 14: GSEA analysis of pathways differentially expressed in MEFs cocultured with macrophages by experimental group.
Supplementary Table 15: Differential gene expression of M1 and M2 vs. M0
Supplementary Table 16: Differential gene expression of M17 vs. M0 and M2
Supplementary Table 17: GSEA of cytokine induced macrophages.
Supplementary Table 18: GSEA of MEFs exposed to cytokine induced macrophages.
Supplementary Table 19: Differential gene expression of MEFS exposed to co-culture CM from M1 and MIL17 vs. M0
Supplementary Table 20: Differential gene expression of MEFS exposed to co-culture CM from M2 and MIL17
Supplementary Table 21: Antibodies used for flow cytometry
Supplementary Table 22: LN biopsies used for IHC and spatial transcriptomics

### METHODS

#### Mice

NZB/W F1 female mice were purchased from The Jackson Laboratory (Bar Harbor, ME) and maintained in a conventional animal housing facility. SLE1.Yaa mice were bred in-house. Due to strain differences, nephritis was defined as fixed proteinuria >100mg/dl for >2 weeks in Sle1.Yaa mice and >300mg/dl for >2 weeks in NZB/W mice. Kidney perfusion, dissociation, and myeloid cell enrichment was performed as previously described. All experimental procedures were conducted in accordance with the guidelines approved by the Institutional Animal Care and Use Committee (IACUC) of the Feinstein Institutes for Medical Research

#### Single-cell RNA sequencing analysis

Human LN biopsies: All stromal cells from the AMP-SLE cohort had undergone QC, analysis, and annotation [1, 2]. Subclustering of fibroblasts and myofibroblasts was performed using Harmony-based batch correction [3]. Using Seurat v5.0, the UMAP was recalculated with a dimension of 30 and a resolution of 0.8. Clustering (FindNeighbors, FindClusters, Seurat v5.0) and differential expression analysis (FindMarkers, Seurat) was used to identify marker genes and manually annotate clusters based on information from the literature [4, 5, 6] and contaminating clusters were removed. ECM Scores were computed by using the function “AddModuleScore" from Seurat (v5.0) at a single cell level and ECM gene sets defined by literature [6]. CNA [7] was performed using histologic activity and chronicity scores from the AMP-SLE dataset. Correlations were filtered by FDR < 0.1. Gene set enrichment analysis (GSEA) of cluster marker genes was performed using fgsea function with ‘biological process’ gene sets obtained from the GO database.

In vitro cocultures: For scRNAseq analysis of cocultured MEFs and renal resident macrophages aligned data was obtained and QC was run using Seurat v5.0. Cells with <2000 features or >7500 features or >7% mitochondrial genes were removed. Datasets were normalized (NormalizeData Function, Seurat), variable features identified (FindVariableFeatures, Seurat, features set to 2000) and scaled (ScaleData, Seurat). Principal components (RunPCA, Seurat) were calculated and plotted as Elbow Plots to individually determine optimal dimensions for UMAP representations. UMAPs were calculated (RunUMAP, Seurat) with dimensions set between 20 and 30 based on previous Elbow Plots and resolution of 1.5. Samples were integrated using a Harmony-based batch correction. For Harmony-integrated datasets the PCA and UMAP representation were calculated based on the Harmony reduction. Downstream analysis was performed as above. The expression of core matrisome genes, provided by the literature [6, 8, 9]were summarized based on normalized gene expression data using the same method as for cell cycle analysis.

**CellChat**

Normalized data were used and cell types and interactions with >10 cells were analyzed. Intercellular communications between each set of 2 cell types were inferred by using the CellChat R package (v2.0). In the ligand-receptor database provided by CellChat [10], paracrine/autocrine signaling interactions (“Secreted signaling”) and extracellular matrix (ECM)-receptor interactions (“ECM-receptor”) were selected for this study. Interaction strength is a measure of the communication probability between a given ligand-receptor interaction and is calculated as the degree of cooperativity/interactions derived by the law of mass action with the expression value of ligands and receptors.

**Flow cytometry**

Isolated single kidney cells were suspended in 1 mL cold PEB buffer (PBS + 2 mM EDTA + 0.5% BSA), followed by incubation with 50 µL magnetic beads (CD45) for 20 minutes on ice in the dark. For magnetic separation, MACS LS columns were used according to manufacturer's' instructions. Eluted cells were suspended in 500 µL PBS/2%BSA containing 1ul Fc block for downstream antibody staining (**Supplementary Table 21**). Staining reactions were conducted in the dark at 4°C for 40 minutes in a 96 well V-bottom plate. Samples were analyzed using a BD FACS Symphony, and cells were sorted using a BD FACSAria. Analysis was performed using FlowJo software.

**Bone Marrow-Derived Macrophages (BMDM)**

BMDM were prepared according to published methods [11]. Using pre-nephritic NZB/W F1 mice, whole bone marrow cells were harvested from the femur and tibia and plated in 6 well plates in 10% FBS complete DME media with 10ng/mL of GM-CSF for 7 days, refreshing the media at day 4. At day 7, the media was replaced with complete media with 10ng/mL of GM-CSF and various cytokines (M0 - Media alone; M1 - 100ng/mL LPS, 20ng/mL IFNγ; M2 - 30ng/mL IL-4, 30ng/mL IL-13; MIL17 - 50ng/mL IL-17a for 6 days; the media was refreshed after 3 days. Six days after stimulation (day 13), BMDMs were washed x 3 and harvested and used for downstream assays and flow cytometry analysis.

**Cocultures**

Primary NZB/W GFP$^{+}$ Mouse Embryonic Fibroblasts (MEFs) were generated from E14.5-15.5 embryos and cryopreserved using previously established methods [12]. Using MEFs at passage 3, 40,000 cells were plated in a 96-well tissue culture-coated plate at Day -1. At Day 0, live, single CD11b$^{+}$ Ly6c$^{-}$ F480$^{+}$ CD81$^{+}$ resident macrophages from nephritic and pre-nephritic NZB/W kidneys were FACS sorted and plated with the MEFs at a ratio of 1:1 in 5% FBS complete DME media. After 48 hours of coculture, MEFs and resident macrophages were harvested for flow cytometry and scRNAseq analysis (10X Genomics) and supernatants retained for scratch wound assays.

**Bulk RNA sequencing analysis**

Total RNA was extracted from cultured cells using RNeasy mini kit (Qiagen). RNA quality was assessed by using an Agilent 2100 Bioanalyzer (Agilent Technologies). Library preparation and RNAseq were performed by Novogene using an Illumina HiSeq 4000, which generated 23–56 million reads per sample. Raw RNA-seq reads were subjected to quality checking and trimming to remove adaptor sequences, contamination, and low-quality reads. The cleaned RNAseq data was then imported into the Partek Flow web platform for downstream analysis. The computational pipeline for expression quantification used STAR aligner. Samples were aligned to mouse transcriptome (mm39) using STAR (version 2.6.1a). Duplicated reads were discovered using Picard tools and removed. Gene annotations were obtained from Ensembl. Good-quality reads were aligned to mouse reference databases, including mm10 mouse genome and genes with a read count <10 across all samples were excluded. R package DESeq2 (v 1.16.1) was used to determine DEGs. Heat maps and volcano plots were generated with heatmap, enhanced volcano plots and ggplot2 R functions, respectively. Significant gene profiles were identified with an adjusted P value of expression differences between groups of <0.05. Gene ontology and signaling pathways were analyzed based on gene profiles using gene set enrichment analysis (GSEA) using the Broad Institutes' GSEA 4.1.0 software. Using 1000 gene set permutations on the Mouse Gene Symbol Remapping Human Orthologs MSigDB.v.7.4.chip Platform, genes were run against three reference gene sets: the KEGG subset of C2 Curated Gene Set, the WP subset of C2 Curated Gene Set, and GO C5 GO Gene Sets. Software and gene lists are freely available at the Broad Institutes' Molecular Signatures database.

**Differential gene expression analysis**

DE analysis, comparing pairs of cell subsets (e.g., myofibroblasts and fibroblasts) was performed using a generalized linear mixed-effects model (GLMM) using the R package glmer. Gene expression was modeled using a negative binomial distribution, with the processing batch, sample collection site and sample ID taken as random effects, and the number of UMIs and percentage of reads mapped to mitochondrial genes included as fixed effects, in addition to the cell subset association.

**Scratch wound**

Scratch wound assays were performed using IncucyteS3 (Sartorius). Primary MEFs were seeded in collagen coated 96-wells (Sartorius) at a density of $4 \times 10^4$ cells/well and incubated for 18 hours in 10% FBS complete DME medium. An open wound area was created in the cell monolayer using the IncuCyte ® Wound Maker tool, and cells were then co-cultured with 48hr conditioned media from strain matched ex-vivo resident macrophage-MEF (NZB/WF1) or polarized BMDM-MEF (Sle1.Yaa) co-cultures. Data was recorded and analyzed using the IncuCyte ® software.

### Spatial transcriptomics

Tissue collection: Human kidney biopsies from patients with lupus nephritis and healthy controls were obtained from Brigham and Women's Hospital with appropriate IRB approval and informed consent (**Supplementary Table 22**). Tissue sections were processed for spatial transcriptomics following the manufacturer's Xenium protocol. Custom probe sets targeting myeloid cluster–specific genes (informed by prior single-cell RNA-sequencing studies [1, 13]) were combined with the manufacturer's multi-tissue panel. Sections were hybridized with probes, imaged on the Xenium platform to capture spatial gene expression, then stained with hematoxylin and eosin (H&E) and re-imaged to align molecular signals with tissue morphology as previously described [2, 13].

Data processing and analysis. Raw Xenium output was processed using Seurat v5.0 and Xenium Explorer v3.1. Initial quality control removed spots/cells in the lowest 10% for unique molecular identifiers (UMIs) and gene counts. Data were normalized, variable features selected, and datasets from multiple kidney sections merged. Downstream processing included scaling, principal component analysis (PCA), uniform manifold approximation and projection (UMAP), and clustering using standard Seurat workflows. To assign cell identities, Xenium data were integrated with published single-cell RNA-seq reference datasets for myeloid cells and fibroblasts [1, 13] using Seurat's anchor-based integration [14]; this produced prediction scores reflecting correspondence to reference cell types while preserving spatial coordinates. Spatial visualization and in situ mapping of myeloid and fibroblast subsets were performed in Xenium Explorer. A board-certified renal pathologist (SS) manually annotated renal compartments to contextualize spatial distributions.

**Immunohistochemistry**

Tissue staining and fluorescent imaging: Five-micrometer FFPE tissue sections from 6 LN biopsies (**Supplementary Table 22**) were mounted on poly-L-lysine-coated glass coverslips. Tissue coverslips were deparaffinized, rehydrated, antigen retrieved by 1xcitrate buffer (pH6, diluted from 100xstock, Abcam ab93678) and then stained with DNA oligonucleotide-conjugated primary antibodies with CODEX staining kit (Akoya Biosciences, 7000008) following their protocol [15]. High-resolution images of full biopsy sections were acquired on an Andor Dragonfly200 Spinning Disk Confocal Microscope. Purified antibodies for the staining included: FAP (clone EPR20021, Abcam, ab271976), aSMA (clone 1A4, Invitrogen, MA1-06110), TREM2 (clone TREM2/7210, NeoBio Technology, 54209-MSM1-P1ABX), FABP4 (clone EPR3579, Abcam, ab219595), GPNMB (clone OTI2F9, Origene, CF807745), MerTK(clone Y323, abcam, ab271851), CD68(clone KP1, eBioscience, #14-0688-82), Claudin1 (clone EPR121871, Abcam, ab238949), and CD163 (clone EDHu-1, Novus Biologicals, NB110-40686). Antibody conjugation was performed using CODEX conjugation kit (Akoya Biosciences 7000009).

**Additional Statistical Analyses**

For comparisons between groups, non-parametric ANOVA with correction for multiple comparisons was used. For analysis of Incucyte data, ANOVA with Friedman's test was used to compare groups that were sampled over time.

**Supplementary Table 1: Demographics of the AMP/SLE Cohort**

| CLASS | All Classes (N = 156) | Class III (N=34) | Class IV (N = 25) | Class V (N = 41) | Class Mixed (N = 56) | Controls (N = 30) |
|---|---|---|---|---|---|---|
| | | | DEMOGRAPHICS | | | |
| Age(mean) | 36.0 | 36.8 | 33.1 | 36.6 | 36.4 | 45.1*** |
| % Female | 86.5 | 91.2 | 96.0 | 78.0 | 85.7 | 76.7 |
| % Hispanic | 28.8 | 26.5 | 32.0 | 22.0 | 33.9 | 3.3*** |
| % White | 31.4 | 38.2 | 24.0 | 24.4 | 35.7 | 80.0*** |
| % Black | 42.9 | 35.3 | 40.0 | 63.4 | 33.9 | 13.3** |
| % Asian | 14.7 | 17.7 | 20.0 | 7.3 | 16.1 | 6.7 |
| | | | HISTOPATHOLOGY | | | |
| | (N = 142) | (N = 31) | (N = 24) | (N = 39) | (N = 48) | |
| Activity Index (range, mean) | 0 – 18, 4.4 | 0 – 14, 4.1 | 0 – 16, 8.4** | 0 - 2, 0.4*** | 0 – 18, 6.0* | |
| Chronicity Index (range, mean) | 0 – 10, 3.8 | 0 - 9, 3.1 | 1 – 9, 4.3 | 0 – 9, 3.5 | 0 – 10, 4.1 | |
| | | | RESPONSE WEEK 52 | | | |
| | (N = 126) | (N = 22) | (N = 21) | (N = 33) | (N = 50) | |
| Complete Responder: | 27.2% | 36.4% | 23.8% | 15.2% | 32.0% | |
| Partial Responder: | 22.4% | 18.2% | 28.8% | 27.3% | 20.0% | |
| Non- Responder | 50.4% | 45.5% | 47.6% | 57.6% | 48.0% | |
| | | | MEDICATION | | | |
| | (N = 154) | (N = 34) | (N = 25) | (N = 41) | (N = 48) | |
| Mycophenolic Acid | 56.1% | 55.9% | 48.0% | 65.9% | 52.7% | |
| Prednisone >10mg | 42.0% | 38.2% | 60.0% | 24.3%* | 49.1% | |
| Cyclophosphamide | 1.3% | 2.9% | 0% | 0% | 1.8% | |
| Belimumab | 4.5% | 5.9% | 12.0% | 2.4% | 1.8% | |
| Hydroxychloroquine | 85.2% | 91.2% | 88.0% | 82.9% | 81.8% | |

**SUPPLEMENTARY FIGURES**

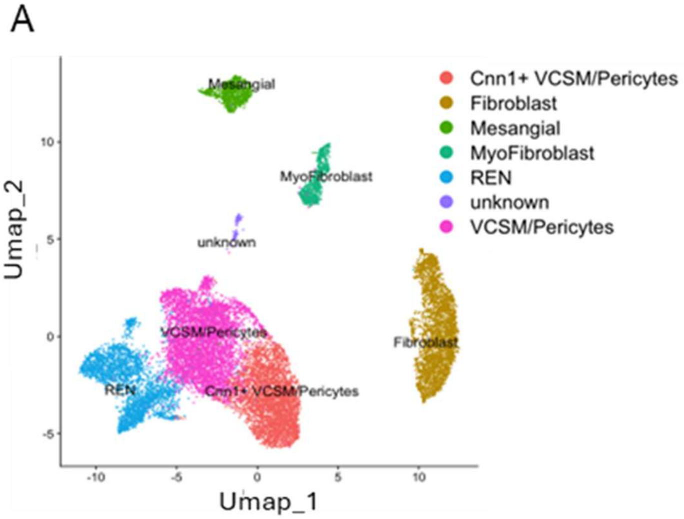


C

| | Control | LN | Total | Control_pct | LN_pct |
|---|---|---|---|---|---|
| Cnn1+ VCSM/Peri1 | 361 | 1149 | 1510 | 21.76 | 7.91 |
| Cnn1+ VCSM/Peri2 | 243 | 1080 | 1323 | 14.65 | 7.43 |
| Cnn1+ VCSM/Peri3 | 5 | 97 | 102 | 0.3 | 0.67 |
| Cnn1+ VCSM/Peri4 | 174 | 608 | 782 | 10.49 | 4.19 |
| Fib1 | 6 | 1052 | 1058 | 0.36 | 7.24 |
| Fib2 | 1 | 1433 | 1434 | 0.06 | 9.86 |
| Fib3 | 2 | 444 | 446 | 0.12 | 3.06 |
| Fib4 | 0 | 92 | 92 | 0 | 0.63 |
| Mesa | 32 | 869 | 901 | 1.93 | 5.98 |
| MyoFib1 | 5 | 462 | 467 | 0.3 | 3.18 |
| MyoFib2 | 8 | 298 | 306 | 0.48 | 2.05 |
| REN1 | 45 | 547 | 592 | 2.71 | 3.77 |
| REN2 | 61 | 743 | 804 | 3.68 | 5.11 |
| REN3 | 131 | 1070 | 1201 | 7.9 | 7.37 |
| REN4 | 3 | 140 | 143 | 0.18 | 0.96 |
| uk | 1 | 133 | 134 | 0.06 | 0.92 |
| VCSM/Peri1 | 45 | 481 | 526 | 2.71 | 3.31 |
| VCSM/Peri2 | 160 | 1133 | 1293 | 9.64 | 7.8 |
| VCSM/Peri3 | 190 | 1368 | 1558 | 11.45 | 9.42 |
| VCSM/Peri4 | 107 | 657 | 764 | 6.45 | 4.52 |
| VCSM/Peri5 | 18 | 211 | 229 | 1.08 | 1.45 |
| VCSM/Peri6 | 61 | 461 | 522 | 3.68 | 3.17 |

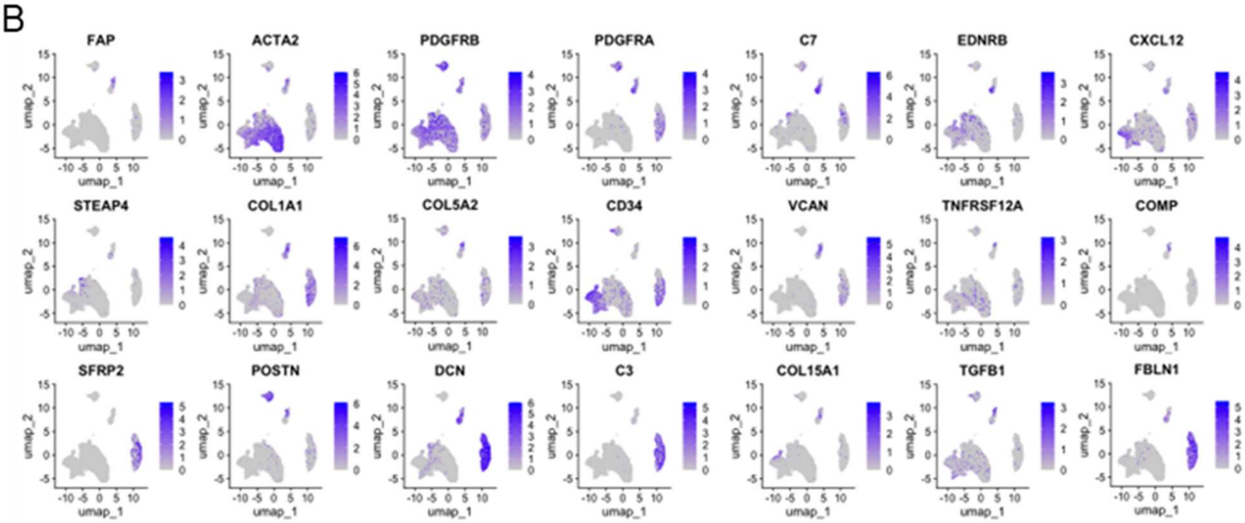


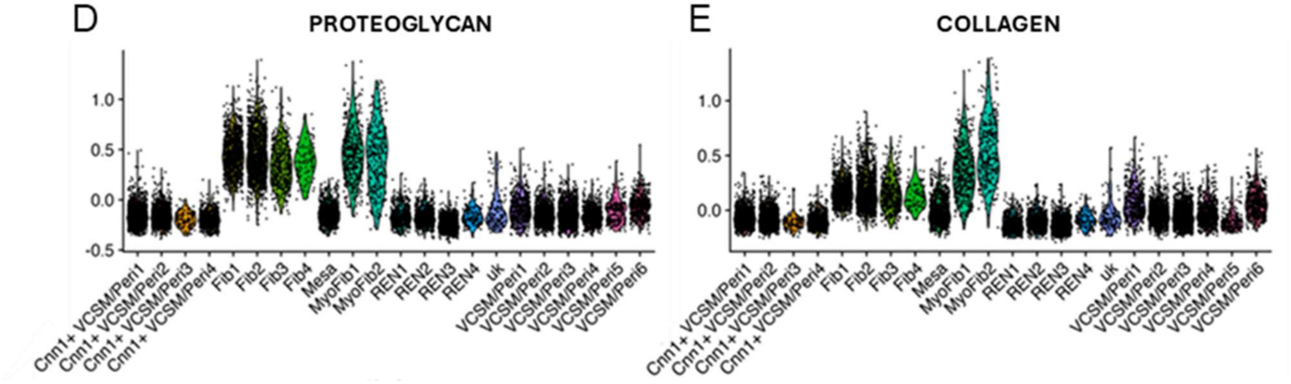

**Supplementary Figure 1:** Stromal cell subsets in LN kidneys. **A-C.** Identification of major stromal cell clusters from 156 LN and 30 healthy donor control kidney biopsies. A. UMAP showing the major clusters of stromal cells. **B.** Feature plots of informative genes in the clusters shown in panel A. **C.** Percent cells in each of the clusters shown in Figure 1A. **D, E**. Violin plots of proteoglycan (D) and collagen gene (E) scores in the clusters shown in C. The small unknown (uk) cluster was removed in Figure 1.

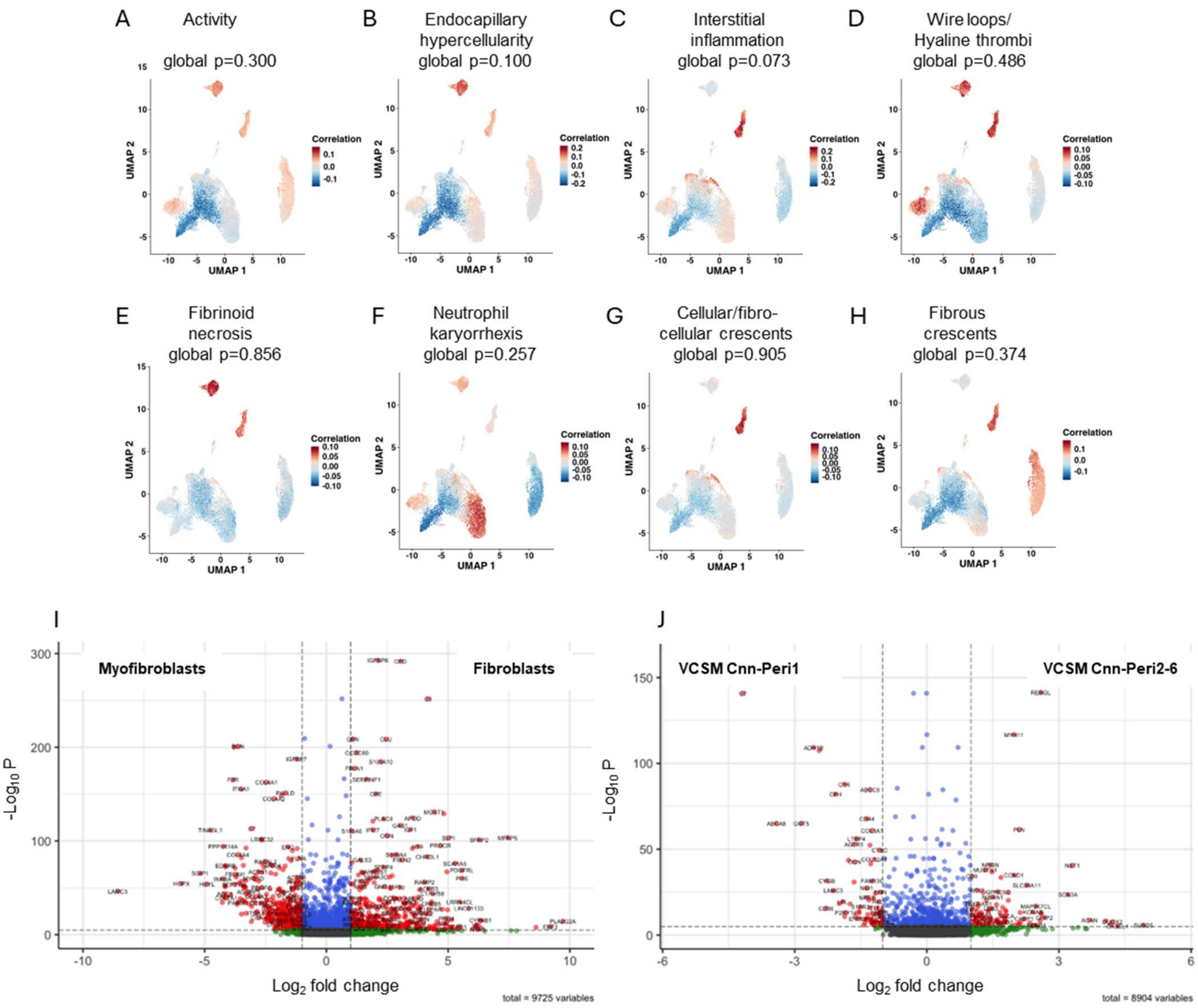


**Supplementary Figure 2:** No association of myofibroblast frequency with disease activity scores in LN**. A-H.** CNA analysis shows no significant association of stromal subset frequency with the total activity score (A) or the indicated subcategories of histologic disease activity and chronicity scores (B-H). **I.** Volcano plot showing differential gene expression of fibroblasts vs. myofibroblasts. **J.** Volcano plot showing differential gene expression of VCSM CNN-/Peri1 vs. the other VCSM CNN-/Peri clusters.

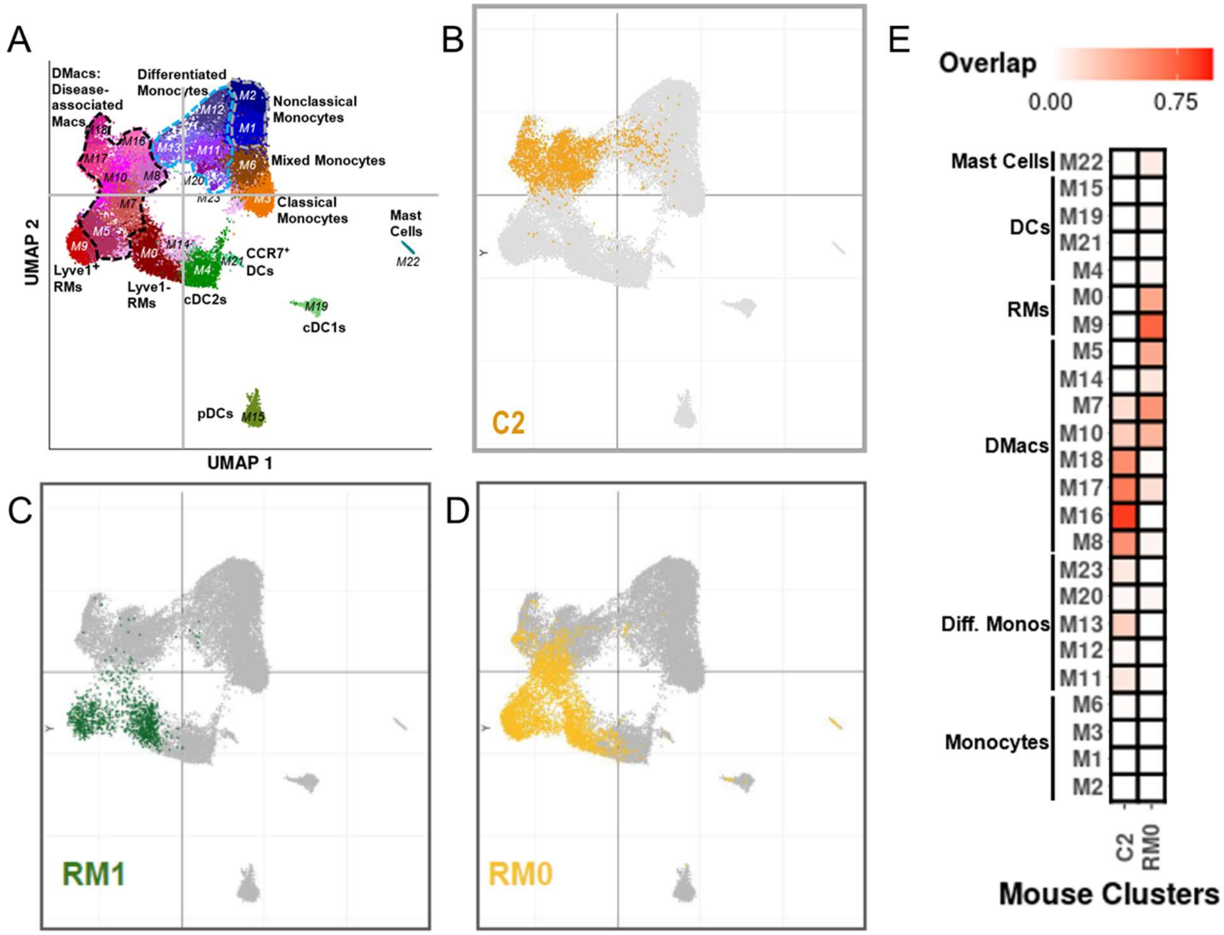


**Supplementary Figure 3**. Classification of myeloid cell subsets in LN kidneys. **A**. UMAP shows fine subclustering of myeloid cells from 156 LN and 30 healthy donor kidney biopsies. Disease-associated macrophage clusters are outlined in black [1, 2]. **B-D**. Mapping of C2, RM1 and RM0 onto the human UMAP. **B**. Orange shows the position of C2 macrophages on the UMAP encompassing clusters M8, 10, 16, 17, 18. **C.** Green shows the position of RM1 macrophages on the UMAP encompassing clusters M0 and M9. **D**. Yellow shows the position of RM0 macrophages on the UMAP encompassing clusters M0, M5 and M9. Clusters M7 and M10 overlap both C2 and RM0. E. Quantitation of the overlap between mouse C2 and RM0 and the 24 human myeloid cell subclusters.

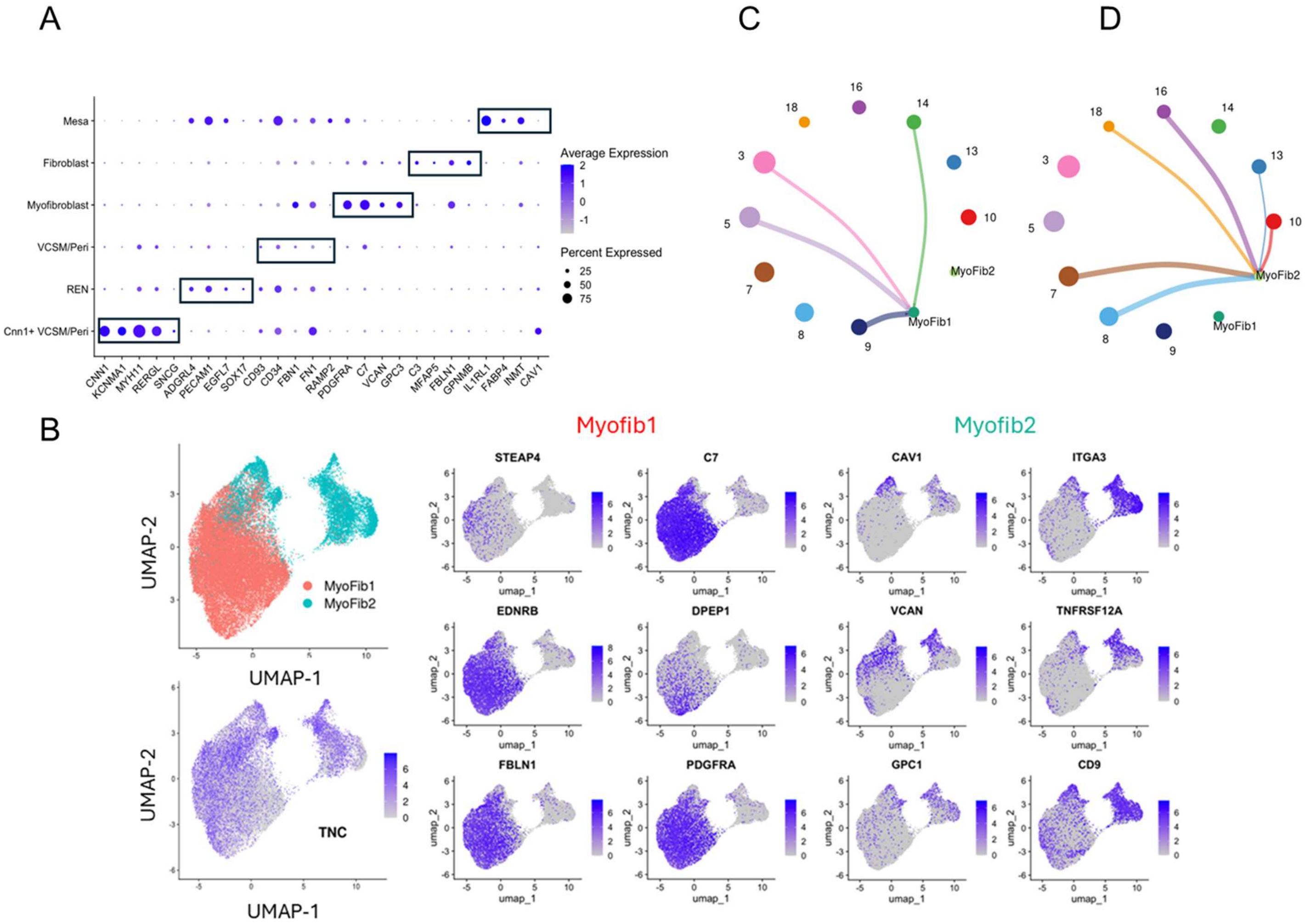


**Supplementary Figure 4**. Identification of stromal cell subclusters in the spatial transcriptomics dataset. **A.** Genes in the spatial transcriptomics dataset that define the major stromal subsets. **B.** Feature plots of myofibroblasts showing genes expressed by Myofib1 and Myofib2. **C, D**. Spider plots showing adjacency of Myofib1 (C)and Myofib2 (D) to macrophage subclusters.

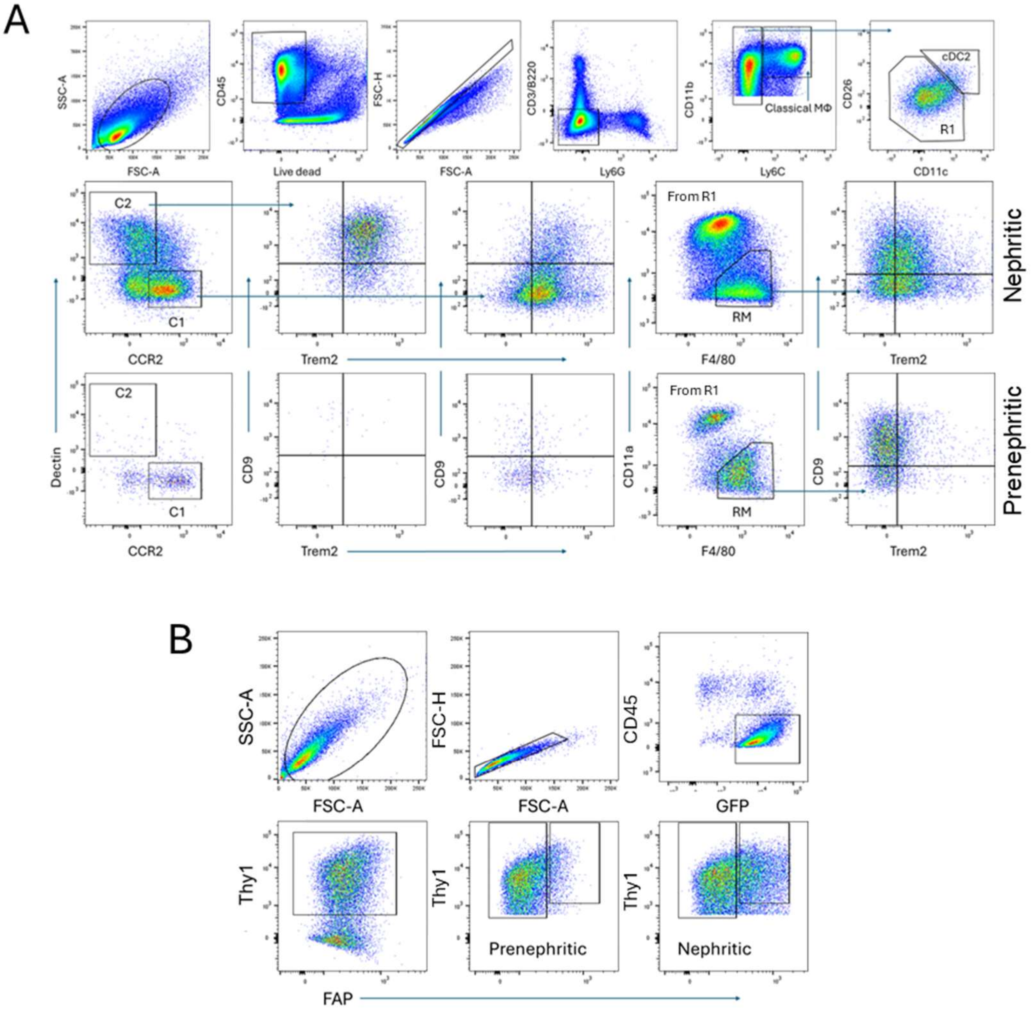


**Supplementary Figure 5**. Flow cytometry gating strategies. **A**. Renal myeloid cell subsets. After gating on live single CD45+CD3-/B220-/Ly6G- cells, CD11bhi cells were selected and gated as Classical monocytes (CD11bhi/Ly6Chi), cDC2 (CD11chi/CD26hi) or resident macrophages (CD11alo/F4/80hi). Classical monocytes were further subdivided into classical 1 (Ccr2hi) and classical 2 (Dectin hi) subsets. Classical 2 monocytes were predominantly found in nephritic mice and were Trem2hi/CD9hi. Resident macrophages from nephritic mice also acquired a Trem2hi/CD9hi phenotype. **B.** Macrophage/fibroblast cocultures. Live single cells from the cocultures were gated for CD45- fibroblasts. The Thy1hi subset was gated to assess expression of FAP.

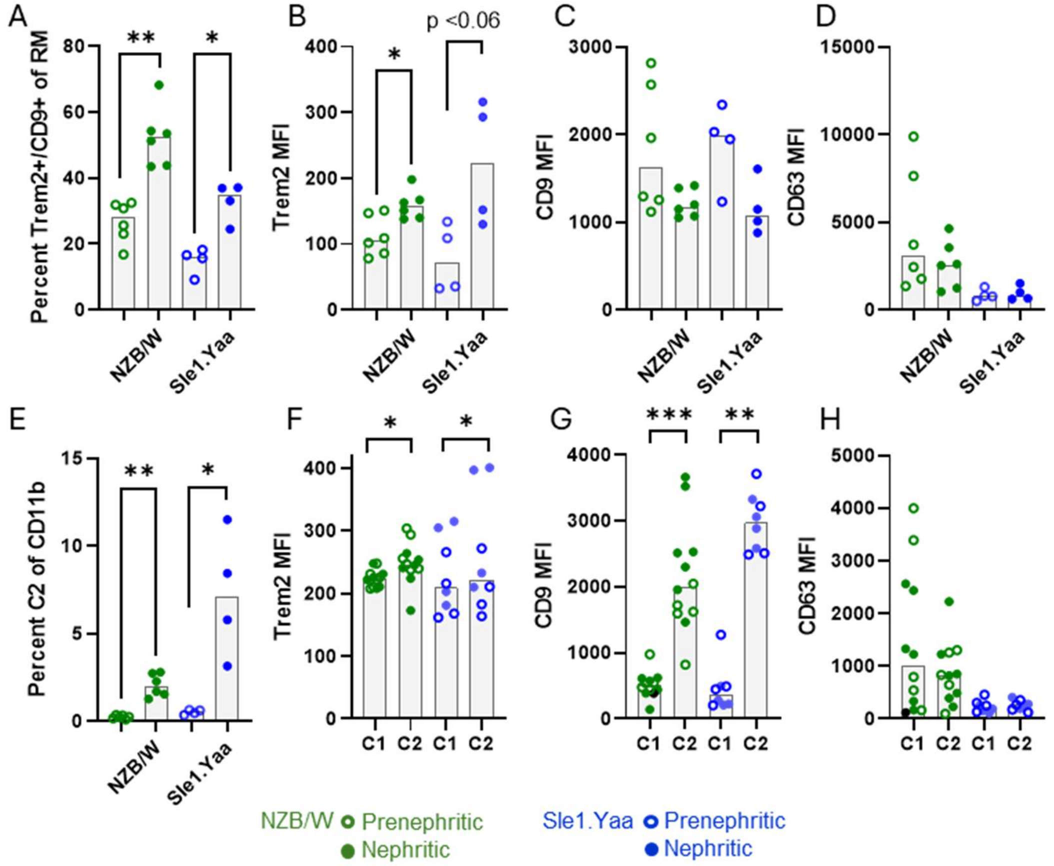


**Supplementary Figure 6**. Enumeration of Trem2hi CD9hi resident macrophage and Classical 2 myeloid cell subsets in kidneys of prenephritic and nephritic NZB/W and Sle1.Yaa mice. **A.** Percentage of Trem2hi CD9hi resident macrophages. **B.** Higher Trem2 MFI in nephritic compared with prenephritic resident macrophages. **C, D.** No change in CD9 (C) or CD63 (D) MFI in resident macrophages between prenephritic and nephritic mice. **E.** Increase in Classical 2 monocytes in nephritic mouse kidneys of both strains. **F, G.** Higher Trem2 (F) and CD9 (G) MFI in Classical 2 compared with Classical 1 monocytes. H. No change in CD63 MFI between Classical 2 compared with Classical 1 monocytes. Kruskal-Wallis ANOVA with Dunn's correction for multiple comparisons. Each symbol represents an individual mouse. *$p<0.05$, **$p<0.01$, ***$p<0.001$.

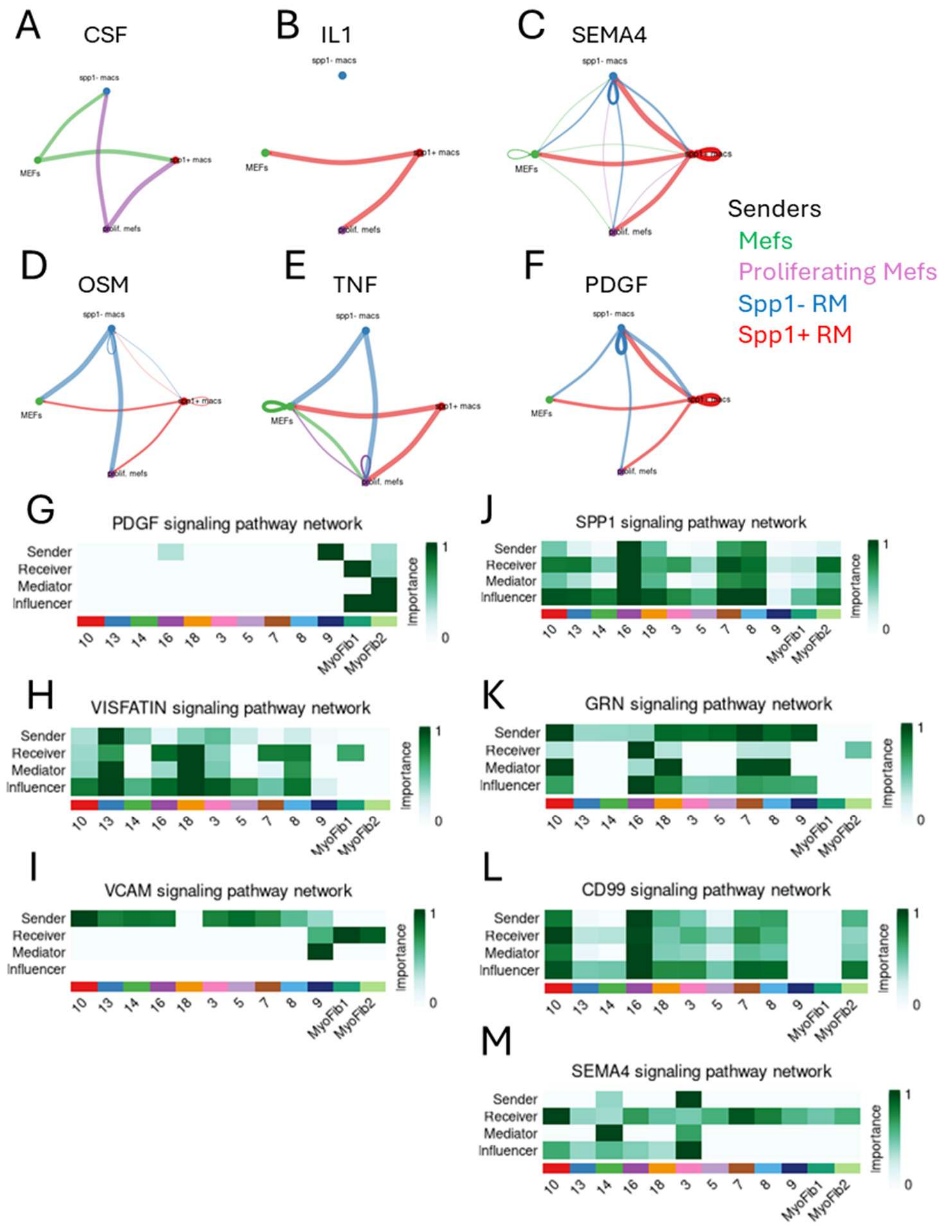


**Supplementary Figure 7**. CellChat analysis of macrophage/fibroblast interactions. A-F. Inferred intercellular communication network for each signaling pathway in the mouse RM/MEF cocultures. Macrophage Spp1+ and Spp1- senders are shown in red and blue lines respectively. Fibroblast senders are indicated by green (differentiated clusters) and purple lines (proliferating cluster). Strength of the potential interactions in the human LN biopsies is shown by the thickness of each line. G-M. Heatmaps showing the relative importance of each cell cluster for each signaling network. G-I show pathways preferentially signaling to Myofib1. J-M show pathways preferentially signaling to Myofib2.

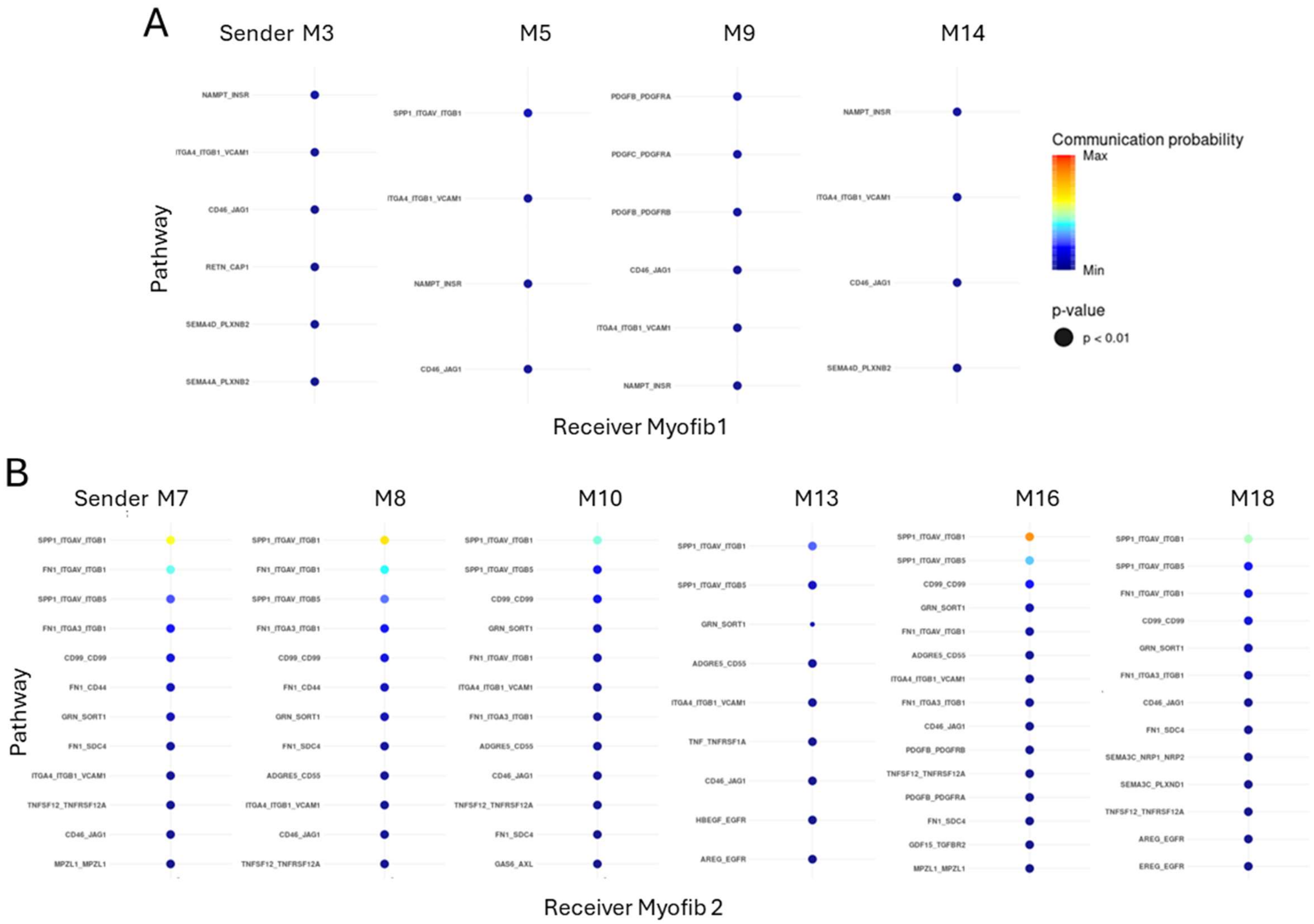


**Supplementary Figure 8**: Potential cell-cell interactions of Myofib1 (A) and Myofib2 (B) with their adjacent macrophage clusters as shown in Figure 2D and 2F indicates frequent interactions between myofibroblasts and macrophages via Spp1.
